# Comprehensive analysis of DNA replication timing in genetic diseases and gene knockouts identifies *MCM10* as a novel regulator of the replication program

**DOI:** 10.1101/2021.09.08.459433

**Authors:** Madison Caballero, Tiffany Ge, Ana Rita Rebelo, Seungmae Seo, Sean Kim, Kayla Brooks, Michael Zuccaro, Radhakrishnan Kanagaraj, Dan Vershkov, Dongsung Kim, Agata Smogorzewska, Marcus Smolka, Nissim Benvenisty, Stephen C West, Dieter Egli, Emily M Mace, Amnon Koren

## Abstract

Cellular proliferation depends on the accurate and timely replication of the genome. Several genetic diseases are caused by mutations in key DNA replication genes; however, it remains unclear whether these genes influence the normal program of DNA replication timing. Similarly, the factors that regulate DNA replication dynamics are poorly understood. To systematically identify *trans*-acting modulators of replication timing, we profiled replication in 184 cell lines from three cell types, encompassing 60 different gene knockouts or genetic diseases. Through a rigorous approach that considers the background variability of replication timing, we concluded that most samples displayed normal replication timing. However, mutations in two genes showed consistently abnormal replication timing. The first gene was *RIF1*, a known modulator of replication timing. The second was *MCM10*, a highly conserved member of the pre-replication complex. *MCM10* mutant cells demonstrated replication timing variability comprising 46% of the genome and at different locations than *RIF1* knockouts. Replication timing alterations in *MCM10*-mutant cells was predominantly comprised of replication initiation defects. Taken together, this study demonstrates the remarkable robustness of the human replication timing program and reveals *MCM10* as a novel modulator of DNA replication timing.

## Introduction

Cell proliferation is one of the most fundamental aspects of development and becomes misregulated in many genetic diseases, in cancer, and during aging and tissue degeneration. A central part of cell proliferation is the replication of DNA, which occurs during S phase and spans roughly a third of the cell cycle in actively dividing cells. Accordingly, delays in DNA replication and S phase completion have been implicated in several developmental diseases that are characterized by growth and developmental defects of various tissues. In addition, some disease-associated gene mutations disrupt the temporal progression of DNA replication, causing certain genomic loci to replicate earlier or later than they normally would (reviewed in (1–3)). Such alterations to replication dynamics have been described in diseases caused by mutations in central DNA replication initiation factors, such as *CDC45* that is part of large deletion occurin in DiGeorge/ Veleocardiofacial (VCF) syndrome (4, 5), *RECQ4* (*RECQL4*) in Rothmund-Thomson syndrome (RTS) (6–9) and components of the ORC and MCM complexes and associated genes in Meier-Gorlin syndrome (10–14). Aberrant replication initiation and progression have also been reported to result from *LMNA* mutations affecting nuclear Lamin A and C in Hutchinson-Gilford progeria (HGPS) (9), *FANCD2* deficiency in a subtype of Fanconi Anemia (FA) (15), *DNMT3B* mutations affecting DNA methylation in ICF1 syndrome (16, 17), and loss-of-function mutations in *BLM* in Bloom syndrome (BLM) (18). These studies used various approaches for assaying replication dynamics, most of which were underpowered to comprehensively characterize the genomic effects of these disease mutations on DNA replication timing. In addition, previous studies haven’t fully considered natural polymorphism in DNA replication timing (19, 20) when interpreting replication timing alterations in disease. Thus, it remains largely unknown to what extent alterations in DNA replication dynamics associate with human developmental diseases. Deciphering these links is important for understanding the etiology of these diseases and for bridging genetic alterations and disease phenotypes via intermediary molecular phenotypes.

Similar to replication timing alterations in developmental diseases, very little is known about the regulatory factors that determine the temporal order of DNA replication progression in mammalian cells. A single well-described modulator of the replication timing program is *RIF1*, which has been shown to regulate global replication timing in yeasts, flies, mice and human cells (21–26). Mutations in *RIF1* cause wide-spread delays and advances in replication timing across numerous regions in the genome, many spanning several megabases of DNA (27–29) and in some cases have even been suggested to define the entire replication timing program (30). Apart from *RIF1*, several studies have described more modest replication timing alterations following knock-out of DNA polymerase theta (Pol θ) (31, 32) or *PREP1* (33). Nonetheless, systematic studies of the effects of *trans*-acting regulators on DNA replication timing are lacking. More generally, the dearth of well-described regulators of DNA replication timing is surprising and warrants further investigation. It could be due to lack of comprehensive assays for testing the effects of *trans*-acting mutations on DNA replication timing, or to a fundamental essentiality of the replication timing program that would preclude the identification of such factors due to cell lethality.

Here, we set to comprehensively test for replication timing alterations in relevant human developmental diseases and in knockouts (KOs) of DNA replication-related genes. We analyzed a total of 184 mutant cell lines and compared them to 167 normal cell lines. Our results point to the rarity of replication timing alterations, suggesting that replication dynamics represent an essential and rigid cellular program. We stress the importance of methodological aspects for the rigorous identification of replication timing alterations and rule out several previously suggested regulators and diseases impacting replication timing. In particular, we identify a set of genomic regions with greater tendency for inter-individual variation, and show that this tendency is independent of genic mutations in *trans*. Last, we report substantial replication timing alterations in *RIF1* KO cells, and - newly discovered here - in a patient carrying mutations in *MCM10*. These results point to the presence of still-elusive factors that regulate human replication timing.

## Results

### Abundant replication timing variation is observed a priori in disease cell lines and gene knockouts

To identify potential modulators of DNA replication timing in human cells, we generated replication timing profiles for 184 cell lines from individuals with genetic diseases or with introduced gene KOs (hereafter, “mutant”) in three cell types, compared to 167 healthy or WT samples (hereafter, “WT”) (**Table 1; Table 2; Supplemental Table 1**). The analyzed cell types included lymphoblastoid cell lines (LCLs), which are EBV-transformed lymphoid cells widely available from many individuals; induced pluripotent stem cells (iPSCs), which are not transformed but not as commonly available across specific patient cohorts; and HAP1 cells, which are nearly-haploid human cell lines derived from a chronic myeloid leukemia patient and readily amenable to CRISPR/Cas9-mediated gene knockout (34, 35). The selected mutant cell lines included those with previous evidence of replication timing alterations, strong links to DNA replication or related pathways such as nucleotide metabolism, DNA repair, chromatin structure (20), and cell lines with alterations in chromosome structure (e.g. disease-associated aneuploidies or repeat expansions). In total, 60 genes or genetic diseases were analyzed across the cell types. DNA from proliferating cell cultures was subjected to whole genome sequencing (WGS) and replication timing was inferred for each sample based on DNA copy number fluctuations along chromosomes, as previously described ((19, 36); see Methods).

**Table 1.** Summary of cell types analyzed in this study. For LCL and iPSC samples, “unique individuals” excludes repeated clones or sequencing of the same individual. In HAP1, all cell lines were derived from a single individual therefore “unique mutant samples” signifies different KO types. HAP1 also includes 26 KO lines involving 22 genes with three genes having multiple KO clones. Among the 26 KO lines, some were sequenced before and after diploidization and are considered the same unique mutant sample. See Supplemental Table 1 for more details.

| Cell type | WT samples | Unique WT individuals | Mutant samples | Unique mutant samples | Unique genetic diseases or genes affected |
| --- | --- | --- | --- | --- | --- |
| LCL | 137 | 124 | 134 | 117 | 32 |
| HAP1 | 6 | 1 | 32 | 26 | 21 |
| iPSC | 24 | 24 | 18 | 11 | 7 |

**Table 2.** Abbreviations of diseases used in this study. Further information is available in **Supplemental Table 1**.

| Disease name | Abbreviation |
| --- | --- |
| Ataxia-Oculomotor Apraxia 2 | AOA2 |
| Ataxia-Telangiectasia | AT |
| Bloom syndrome | BLM |
| Breast Cancer, Type 1 | BRCA1 |
| Beckwith-Wiedemann Syndrome | BWS |
| Cornelia de Lange Syndrome 1 | CDLS1 |
| Chronic myeloid leukemia | CML |
| DiGeorge Syndrome | DGS |
| Myotonic dystrophy type I | DM1 |
| Fanconi anemia | FA |
| Friedreich's ataxia | FRDA |
| Fragile-X syndrome | FXS |
| Huntington's disease | HD |
| Hutchinson-Gilford Progeria Syndrome | HGPS |
| Immunodeficiency-centromeric instability-facial anomalies syndrome 1 | ICF1 |
| Lesch-Nyhan Syndrome | LNS |
| Mental retardation, autosomal dominant 1 caused by a deletion in MBD5 | MRD1 |
| Rothmund-Thomson Syndrome | RTS |
| Rett Syndrome | RTT |
| Rett Syndrome, congenital variant | RTTC |
| Spinal and bulbar muscular atrophy | SBMA |
| Spinocerebellar ataxia type 1 | SCA1 |
| Seckel Syndrome | SCKL1 |
| Sotos Syndrome 1 | SOTOS1 |
| Down Syndrome | TRI21 |
| Williams-Beuren Syndrome | WBS |
| Wolf-Hirschhorn Syndrome | WHS |
| Werner Syndrome | WRN |
| Turner Syndrome | XO |
| Translocation of the X chromosome | XTRANS |
| 4 X chromosomes | XXXX / XXXXY |
| Klinefelter Syndrome | XXY |

Replication timing profiles had a median Pearson’s correlation coefficient of 0.86 (0.31-0.90) among all LCL samples, 0.86 (0.71-0.90) among LCL WT samples, and 0.90 (0.73-0.95) among LCL WT repeat samples (**Fig1A-B**, **Fig S1**). There was also a high correlation (r = 0.94) of the WT LCL sample NA12878 to its replication profile generated by sequencing S and G1 phase DNA (37), further demonstrating the high quality of replication timing profiles generated in this study (**Fig 1C**). For iPSCs and HAP1 cells, the median between-sample correlations were 0.95 (0.66-0.96) and 0.90 (0.66-0.93), respectively. The correlations of samples within a given cell type were somewhat lower than expected, which we attribute to several low-correlating mutant samples (to be further discussed below), a less stringent approach to filtering in anticipation of replication timing variation (see Methods), and the variability of sample source and WGS method and sequencing depth.

**Fig 1.**
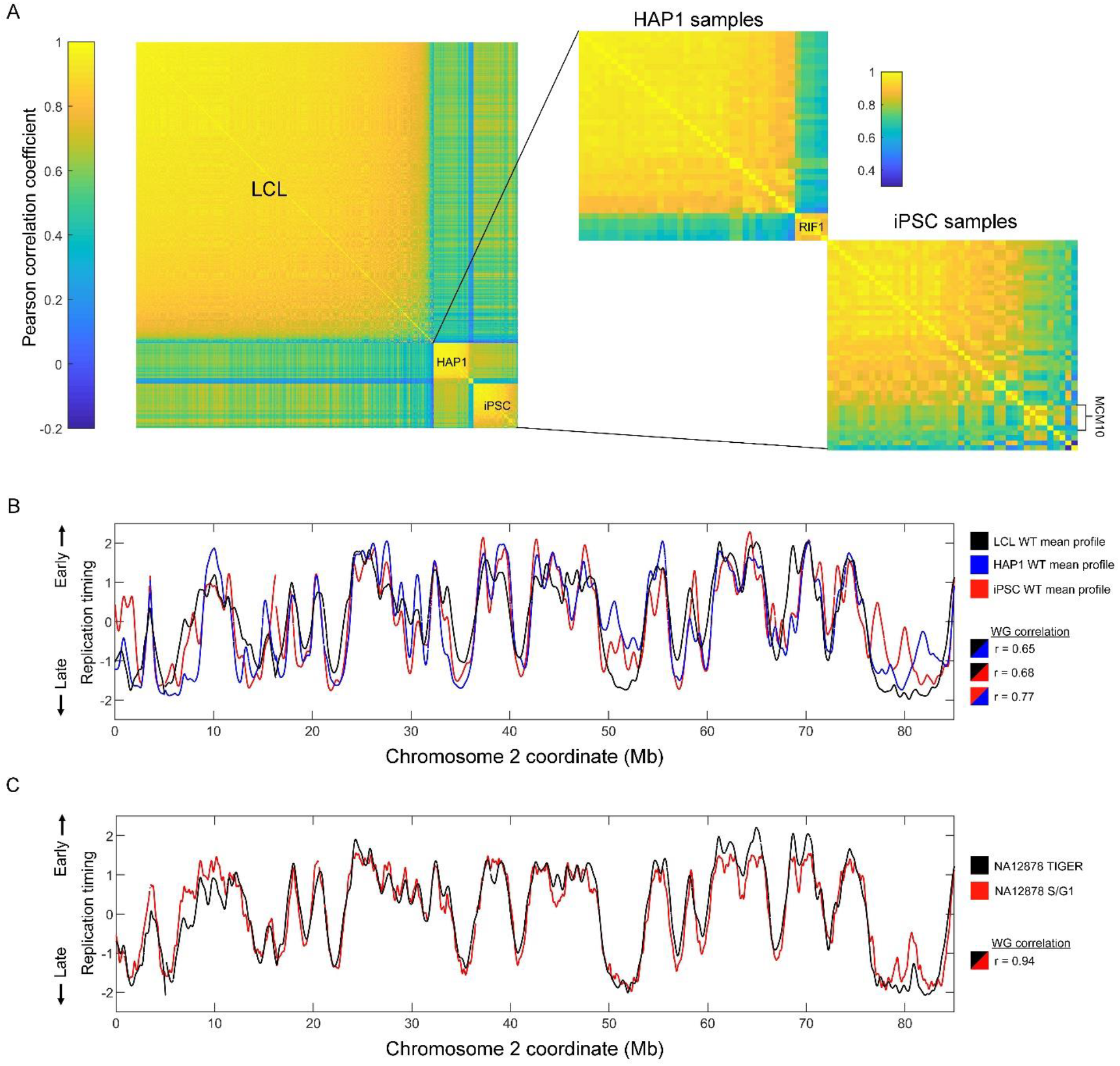
Overview of replication timing data. (A) Correlation matrices of replication profiles for all samples. (B) Replication profiles and whole genome correlations of the WT mean profiles of the different cell types. (C) Replication timing profile comparison and whole genome correlation for the sample NA12878 generated with TIGER (see Methods) or S/G1 sequencing.

To analyze replication timing variation between mutant and WT cell lines, we performed an analysis of variance (ANOVA) in sliding windows across the autosomes (sex chromosomes were not considered since the samples included both male and female individuals). ANOVA was applied to raw data (before smoothing) in windows of 185kb of uniquely alignable sequence (76 bins of 2500bp; Methods) with a step of a quarter window. We studied both individual samples as well as samples grouped by mutated gene or genetic disease and compared each to the control samples of the same cell type. Individual windows with a Bonferroni-corrected p-value <0.05 were considered to have variant replication timing, and overlapping variant windows were subsequently merged into continuous variant regions. Given the variable numbers of mutant and WT samples in the grouped analyses, we permissively allowed samples to show an inconsistent direction of replication timing difference compared to other samples, as long as the group ANOVA was significant.

All samples grouped by mutated gene, and nearly all individual mutant samples (177/184), contained at least one genomic region with *a priori* replication timing variation compared to the WT samples (**Fig 2A, Fig S2A**). Across all cell types, individual mutant samples showed a median variant replication timing covering 1.33% of the autosomes. Samples grouped by mutated gene showed a median replication timing variation covering 6.67% of the autosomes. *RIF1* KOs in HAP1 cells contained the highest proportion of variant replication timing at 72.65%, with the five individual samples ranging from 43.79% to 71.47% genome variation compared to WT (**FigS2B**). Several other gene mutations were also associated with higher-than-average genome variation (e.g., RTT, RTS, *MCM10*, FXS in iPSCs).

**Fig 2.**
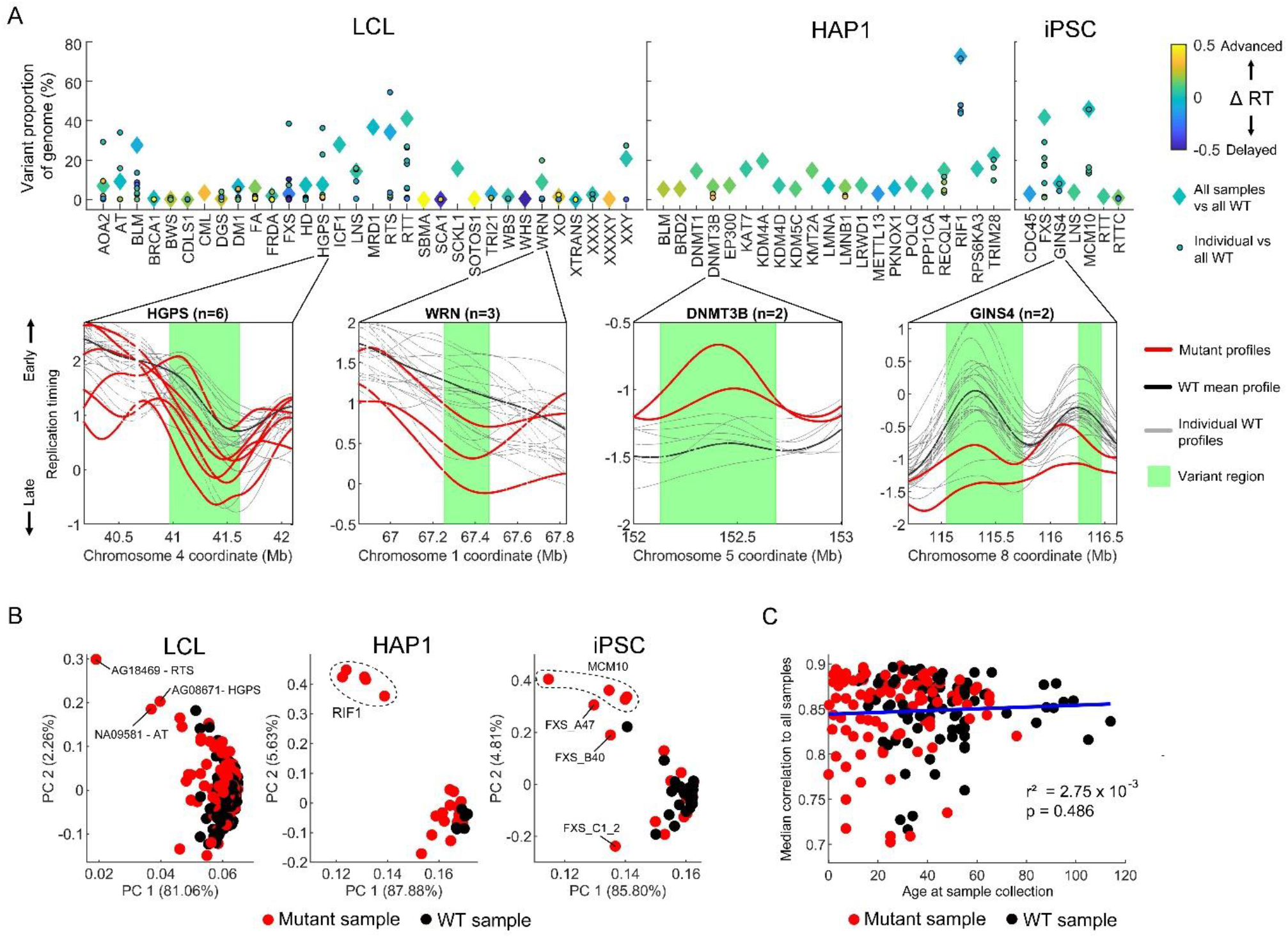
Analysis of variance detects significant replication timing variation in all mutated gene groups and most individual samples. (A) Top: Proportion of the autosomal genome with variant replication timing detected via ANOVA with a Bonferroni-corrected p-value of <0.05. For each gene mutation, both grouped and individual samples were evaluated against WT samples for each cell type. Colors represent the extent of bias towards replication timing advances or delays. Bottom: four example gene mutations are shown, with variant regions depicted based on the grouped analysis. For LCL samples, only 20 WT samples are shown. (B) PCA of the entire autosomal replication timing profiles. Clustering by mutation is observed for the *RIF1* KO in HAP1 cells and for *MCM10* mutants in iPSCs. (C) Relationship of individual donor age at sample collection to median correlation of autosomal replication timing among LCL samples. Only samples with a mean correlation >0.7 to other LCL samples are included (see Methods and Fig S3B for all samples).

In the 99,850 variant regions called across all mutated gene groups and individuals, the median absolute difference in replication timing from the mean WT value was 0.50 units of standard deviation. In virtually all of the 30 strongest cases (with variation spanning ≥20% of the genome), replication timing variations included both advances and delays in roughly equal proportions; the most directionally-biased sample was a HAP1 *RIF1* KO sample with a mean delay across variant regions of 0.14 standard deviations (**Fig 2A**).

Despite many mutated gene groups showing a substantial genomic proportion of variant replication timing, closer inspection ruled out most as candidates for gene-related dysregulation of replication timing. For example, replication timing in variant regions for AOA2, RTS, BLM, HGPS, and FXS (in LCLs, across autosomes) were largely driven by one or two outlier samples while the rest strongly resembled the WT profile (**Fig S2C**). Accordingly, principal components analysis (PCA) of replication timing profiles did not reveal clustering by mutated gene except for *RIF1* KO in HAP1 cells and *MCM10* mutants in iPSCs (**Fig 2B**). Singular outlier samples with abnormal replication timing will be further investigated below.

Apart from disease state, we also assessed other factors that could influence local or global replication timing variation in our samples. Sequencing cohort or batch effects seemed to be minimal based on PC analysis, with the exception of one cohort of samples, which had the lowest-coverage sequencing (**Fig S3A**). We also ruled out that replication timing is significantly influenced by a person’s age in our sample set. First, we compared the correlations of replication timing profiles in 186 LCLs from donors of known ages (ranging from 0 to 114 years old) at sample collection (**Fig 2C, S3B**). If replication timing changes with age, we would expect deviation from the average replication profile in older (or younger) samples. However, we found no change in replication timing correlation to other samples as a function of a sample’s age. Secondly, PCA of WT replication timing profiles did not show stratification by age (**Fig S3C**). Although the unknown number of cell culture passages that each cell line underwent may confound the analysis of ‘age’, the large number of samples analyzed here effectively rules out a strong influence of aging on replication timing, at least in LCLs. Also consistent with the minimal or no effect of age on LCL replication timing, we did not observe any notable replication timing variability in diseases associated with accelerated aging (HGPS, WRN, BLM) (**Fig 2A**, see further below).

Taken together, variability in replication timing is detected in most mutant individuals and mutated gene groups. However, variability is not necessarily the result of a gene mutation-related modulation of replication timing, but may instead be driven by a subset of outlier samples or by background technical or biological variation, as further explored below.

### Recalibration of false discovery rates using simulations of replication timing variation

Identifying genuine variability in replication timing as a result of a diseased state or gene mutation may be confounded by background replication timing variability, which may arise due to technical factors or be related to common polymorphisms that influence local replication timing (19, 20). For example, as shown above, a subset of outlier samples led to the identification of variability in replication timing that is not shared with other samples of the same disease or gene mutation. Variant detection can be made stricter by adjusting the significance threshold or by requiring that all samples within a group follow the same trend. These remedies are expected to be heavily influenced by the number of samples compared in each mutant gene and WT group and were therefore not implemented in initial analyses.

When inspecting the ANOVA variant results using quantile-quantile (QQ) plots, we observed widespread inflation (and in some cases, deflation) in the obtained p-values (**Fig 3A, Fig S4A**). This inflation is likely related to the continuous nature of the DNA replication profiles from which the data is sampled. It did not result from the sliding window method, as it was still observed in an ANOVA variant search with non-overlapping windows (**Fig S4B**). Importantly, the extent of p-value inflation was different for each mutated gene group and individual, which makes it challenging to determine an appropriate threshold for rejecting the null hypothesis. Strict multiple testing correction did not mitigate this challenge, as the p-values were inflated beyond Bonferroni-corrected significance thresholds. An alternative method for multiple-test correction that could be considered in this case is q-value transformation, where p-values are adjusted based on false-discovery rate (FDR). However, in the ANOVA tests for replication timing variation, such FDR would be independently calculated based on the p-values for each mutant group or individual. This creates a different FDR value for each analysis even when the number of mutant samples being analyzed is equal and the WT samples remain the same (**Fig S4C**).

**Fig 3.**
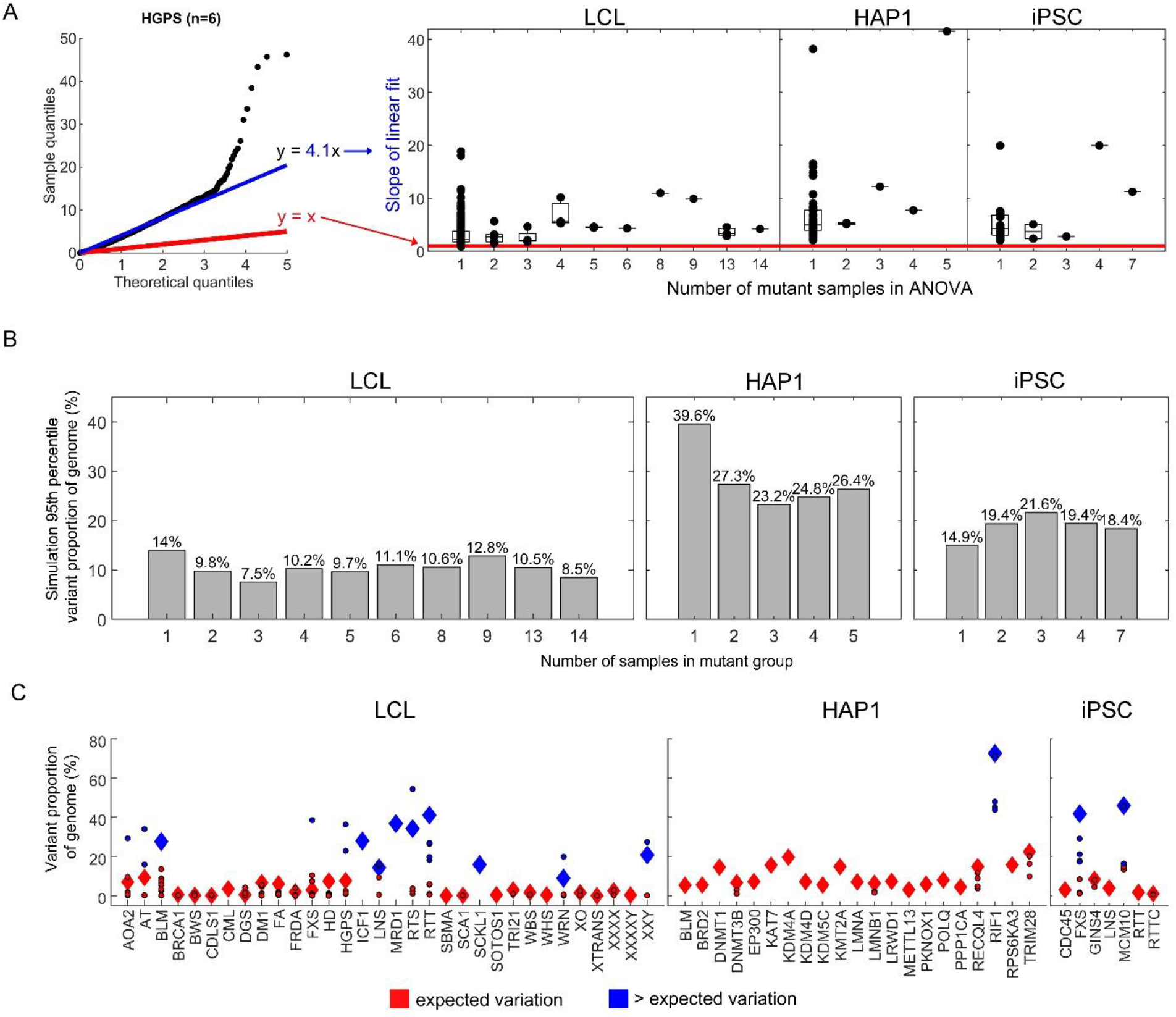
Variability in replication timing. (A) p-value inflation quantified with QQ plots for the different mutant-WT ANOVA tests. Theoretical quantiles (the uniform distribution of p-values) and ANOVA test quantiles should be linearly related with a slope of 1 (red) if they are generated from the same distribution. The linear fit of the ANOVA quantiles to theoretical values (blue) quantifies the deviation as inflation or deflation of p-values. The boxplot demonstrates the p-value inflation statistic in the different ANOVA tests for mutated gene groups and individuals. (B) The 95^th^ percentile of variant proportions of the genome from the 1000 simulations for different mutant group sizes. The number of WT samples matched that available for each cell type (137 in LCL, 6 in HAP1, and 24 in iPSCs). (C) Same as Fig 1A, highlighting (blue) the mutant groups and individuals exceeding the 95^th^ percentile cutoff.

To overcome these statistical challenges of analyzing sample groups with different sizes, we sought to determine a universal FDR by empirically calculating the expected variability in our replication timing data for any relevant number of compared samples. To achieve this, we permuted the samples, randomly assigning them into mutant and WT groups and repeating the ANOVA variant search. Given that the majority of mutant samples fell within the correlation distribution of WT samples (**Fig 1A**), we used all samples (WT and mutant) with a correlation >0.7 to other samples. This cutoff removed genuine outlier samples such as *RIF1* and others, while the inclusion of mutant samples in the “WT” permutations maximized the number of samples available for this analysis. The latter enabled us to test permuted WT groupings up to the actual number of WT samples available for a given cell type. We thus tested WT and mutant groups with varying numbers of samples in each, performing 1000 sample permutations for each pair of groups sizes. In this analysis, FDR is akin to the expected variability of replication timing based on the number of samples in each compared group.

In these “simulations” of background variability, the proportion of the genome found to be variant decreased with increasing numbers of WT samples, for all cell types (**Fig S5**). This is expected, since including more controls effectively rules out many false positive outlier observations. Importantly, there was a substantial dispersion around the median variation across the simulations (**Fig S6**). For example, in simulations of LCLs with one mutant sample and 137 WT samples (i.e. all available), the median genome variation was 0.18% but the 5^th^ and 95^th^ percentile (representing typical limits of low and high variation) were 0% and 13.97%, respectively. Therefore, we would argue that 13.97% genome variation (the 95^th^ percentile) represents an upper limit for expected variation in analyses with one mutant and 137 WT samples. To further illustrate the application of a 95^th^ percentile cutoff for expected variation, consider simulations for LCLs with three mutant and 137 WT samples – the same number of actual samples available for *BRCA1*, DGS, SCA1, and WRN. In this case, the median proportion of the genome with replication timing variation was 0.28% and the 95^th^ percentile was 7.54%. If we use the 95^th^ percentile as the upper cutoff for expected variation, then among the LCL mutant groups that had three mutant samples we would rule out *BRCA1* (0.58%), DGS (0.51%), and SCA1 (0.15%) as having an extent of variation within the expected range (**Fig 3B**). In contrast, the group containing the three WRN samples does have an extent of variation (8.93%) exceeding this 95^th^ percentile cutoff and would therefore remain as a candidate for later analyses of replication timing variation.

We applied the 95^th^ percentile cutoff to all samples and groups (**Fig 3B**), which resulted in 155 of 184 individual mutant samples, and 52 of 60 mutated gene groups being classified as within the expected range of replication timing variability (**Fig 3C**). Therefore, these gene mutations (at least in the analyzed cell type) were concluded to likely not influence replication timing. The several mutants that did exhibit variation above the expected range of replication timing variability will be analyzed further below. Background heterogeneity in replication timing data still emerges as a critical factor requiring rigorous consideration in any search for replication timing differences between samples or groups.

### Replication timing variation is non-randomly distributed across the genome

Based on the above simulations, variability in replication timing can be expected in a substantial percentage of the genome, depending on cell type. We asked if this variability (in both simulations and true analyses) is uniformly distributed across the genome or clustered in specific regions and/or specific replication times. For the true mutant individuals and gene groups, the regions of variant replication timing were bimodally distributed to late- and early-replicating regions of the genome (**Fig S7**). Furthermore, variant regions across all individual mutant analyses overlapped more than would be expected by chance. The median proportion of variant region coordinates shared between two mutant samples of the same cell type (excluding samples without variant regions) was 2.43% (**Fig S8A**). Comparatively, variant regions only covered a median of 1.33% of the autosomes in individual mutant analyses so therefore we would roughly expect only 0.0177% of coordinates to overlap by chance (2.43% x 2.43%). Surprisingly, variant regions across different cell types also showed high overlap (**Fig S8B**). Using a Fisher’s exact test, the average p-value for variant region overlap across all mutant samples (including analyses where mutant sample pairs did not overlap at all) was 3.77 x 10^-5^ (**FigS8C**; 7.47 x 10^-8^ among mutant sample pairs with non-zero overlap). Taken together, we conclude that replication timing variability tends to localize to particular regions of the genome.

To analyze where replication timing variation tends to occur in the genome in each cell type, we used the randomized sample grouping simulations to calculate the median variation p-value in each sliding window across the genome. Replication timing variation was disproportionately more common in early- and late-replicating parts of the genome (**Fig S9**) in a similar bimodal distribution to where true mutant variant regions fell (**Fig S7**). Notably, variation appeared to be greatest in the earliest replicating regions of the genome, suggesting that zones of DNA replication initiation tend to be more variable between samples. Indeed, greater replication timing variation was often observed at peaks, as well as valleys, in the replication timing profiles, which represent regions that contain sites of DNA replication initiation and termination, respectively (**Fig 4A**). Notably, different peaks and valleys showed different intensities of variation. The genomically-variable regions detected in the simulations were consistent with the variable regions in the analyses of mutant samples (**Fig S10**).

**Fig 4.**
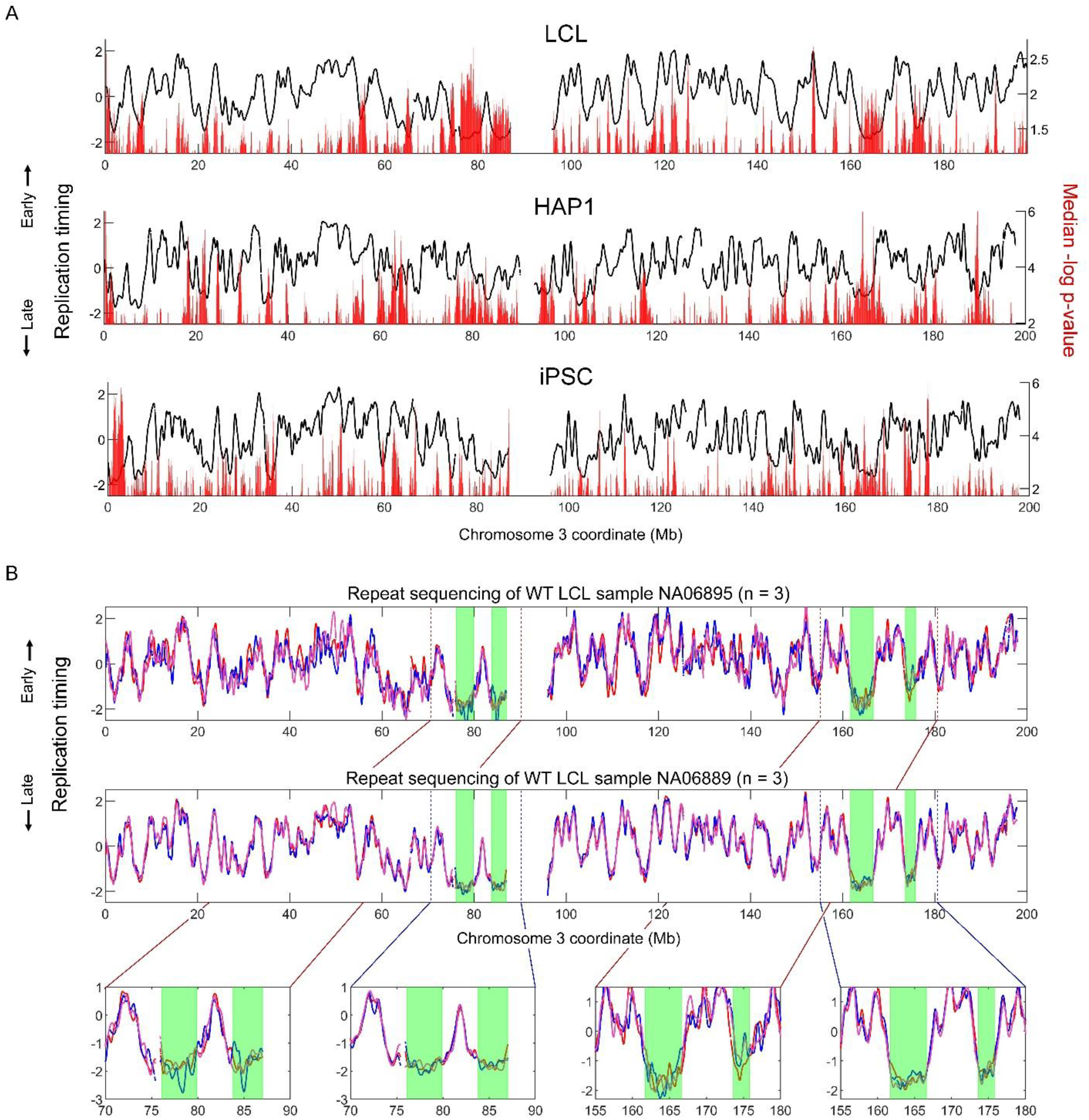
Localized variability in replication timing. (A) The median p-value in each sliding window across 1000 simulations using all mutant group sizes tested against the total number of WT samples available for each cell type (137 for LCL, 6 for HAP1, and 24 for iPSCs). Only p-values below the median are shown. (B) Replication timing of the LCL WT samples NA06895 and NA06889 (each of which includes three repetitions). Late-replicating regions with notable variation are highlighted in green.

Another notable category of replication timing variants were large (>1Mb), very late-replicating regions void of clearly defined peaks (e.g., **Fig 4A-B**). These structures were most prominently present in LCLs. For example, chromosome 3 in LCLs contained four of these late-replicating regions that together covered 14Mb (**Fig 4B**). Within these regions, even replication timing of repeat samples from the same individual varied considerably. For example, in three repetitions each of the WT LCL samples NA06895 and NA06889, the correlation of replication timing within these late-replicating zones was on average 0.69 and 0.65, respectively, markedly lower than the correlations of 0.89 and 0.97 for the rest of the chromosome. Taken together, some genomic regions are relatively enriched for background variation and could be expected to show up in any sample comparison, whether mutants compared to WT or in control comparisons.

### Most samples with high variability in replication timing are false-positives

By defining the range of background replication timing variability, 29 individual mutant samples and 9 mutated gene groups were identified as candidates for representing *trans*-acting modulators of replication timing (**Fig 3C**). Of those, three candidate LCL mutated gene groups – ICF1, MRD1, and SCKL1 – contained only one sample each. Despite having greater than expected variation compared to WT samples, MRD1 and SCKL1 had an overall high correlation to the mean LCL WT replication timing profile (r = 0.85 and 0.92, respectively). We suspect that these samples may show a high degree of variation due to low frequency subclonal deletions or duplications that might have escaped filtering and were ultimately amplified during data processing (specifically, normalization of the genome to a copy number of two before the ANOVA variant search; see Methods). Therefore, we do not consider individual samples with both high replication timing variability and high correlation to WT samples as strong candidates for having altered replication timing. In contrast, ICF1 showed both high variability and low correlation (r = 0.67) to the mean LCL WT replication timing profile. Since we only analyzed a single sample with this disease in LCLs, we cannot rule out other explanations for the high level of variation in this sample (e.g. a secondary somatic mutation, or technical factors). Furthermore, we analyzed two repeats of a HAP1 KO of the DNA methyltransferase *DNMT3B* gene, mutated in ICF1 syndrome, but did not find elevated replication timing variability. We thus conclude that ICF1/*DNMT3B* is not a strong candidate for altered replication timing.

Following a similar rationale, we eliminated additional individual mutant samples with high replication timing variability yet high correlation to the mean WT profile. In LCLs, 15 individual mutant samples exhibited replication timing variation above expectation (**Fig 3C**), of which seven demonstrated high correlation (r > 0.8) to the mean LCL WT profile (**Fig S11**). These included sole individual outliers among WRN and XXY samples, effectively removing this gene mutation and aneuploidy state, respectively, as candidate regulators of replication timing. Among the eight LCL individual mutants with low (r ≤ 0.8) correlation, six were from the cohort with the lowest-coverage sequencing. Given this overrepresentation of low-coverage samples, we regarded these samples as possible false positives; this further eliminated the RTS sample group as an *a priori* candidate for regulating replication timing, as well as the gene mutations in AT, HGPS, LNS, and RTT – all of which contained outlier samples that belonged to either this low-coverage or to the high-correlation samples. This left two remaining individual mutant outliers, in AOA2 and in FXS. However, re-sequencing these individual samples did not reproduce the high replication timing variability, ruling out the corresponding gene mutations as likely causes of the initially observed variation. No gene mutations were eliminated in HAP1 cells nor iPSCs based on individual mutant sample correlation to the mean WT profile.

After the above elimination, four mutated gene groups remained. The mutated gene group of Fragile-X syndrome (FXS) in iPSCs (although not in LCLs), and four of its seven individual mutant samples, had replication timing variability above the expected background threshold. Of the FXS iPSC individual mutant samples, three showed low correlation to the WT mean profile (**Fig S12**). However, the abnormal replication timing was not shared among the clones or resequenced samples available for two of the three FXS mutant individuals. Therefore, based on the available samples we conclude that the *FMR1* gene, mutated in FXS, is unlikely to be a *trans*-acting regulator of replication timing as the associated variation is not consistently observed among genetically identical samples.

There were nine mutant samples from four individuals with Bloom syndrome (BLM), which showed variant replication timing covering 0.14% to 13.6% of the genome and 27.6% when analyzed as a group (**Fig 2A**). Of those, two individuals (NA04408 and NA09960) as well as their re-sequenced samples showed high correlation to the mean LCL WT replication timing profile and relatively invariant replication timing (**Fig S13**). The remaining two individuals (NA03403 and GM16375) as well as their re-sequenced samples had significant replication timing differences from WT samples, typically encompassing novel or lost peaks (**Fig S13**). These peak gains and losses as well as the overall replication profiles were not fully consistent among the three experimental repetitions of sample NA03403, suggesting possible technical or biological noise in this particular individual. While we hypothesize that true replication timing variation is present in these two BLM samples, we refrain from ascribing them to the *BLM* gene mutation directly given the lack of consistency across all BLM samples. Moreover, a *BLM* KO in HAP1 failed to show altered replication timing, further suggesting that the *BLM* gene is not directly involved in global replication timing regulation. It is possible that the replication timing variability observed in only half of the LCL BLM individuals may occur due to a secondary somatic mutation in another, potentially unknown regulator of DNA replication timing, especially considering that loss-of-function mutations in the BLM RecQ helicase results in increased somatic crossing-over and spontaneous mutation rate (38).

### MCM10 is a novel regulator of DNA replication timing

After a systematic analysis of 60 mutated genes or genetic diseases, only mutations in *MCM10* and *RIF1* demonstrated consistent variability in replication timing, low correlation to WT replication timing, and clustering of replication timing in PCA all consistently related to the mutated gene (**Fig 2A**, **Fig S14, Fig 2B**). *RIF1*, a known modulator of replication timing, showed variant replication timing covering 72.65% of the genome. The locations of variation were shared among the individual HAP1 *RIF1* mutant samples, which also had highly correlated replication timing profiles at a similar level to the correlation of WT HAP1 samples (**Fig S14**).

*MCM10* mutant cell lines demonstrated high deviation in replication timing from the corresponding WT iPSC profile, with variation covering 46.0% of the genome. We verified that the *MCM10* iPSCs were not spontaneously differentiated (and therefore demonstrating the replication timing of another cell type) by comparing them to various repli-seq profiles of differentiated cells (39) (**Fig S15**). We also confirmed that the abnormal replication timing in *MCM10* samples was not the result of copy number alterations that escaped filtering (**Fig S16**).

Given the established and unique role of *RIF1* in DNA replication timing, it is possible that the *MCM10* mutations operate in the same pathway or indirectly (e.g. by means of a secondary mutation) impinge on *RIF1* function. To compare the *MCM10* profiles to *RIF1* profiles in a similar cell type, we generated two *RIF1* KO clones in haploid ESCs (40, 41) using CRISRP/Cas9. The WT ESC and iPSC replication profiles were similar (**Fig 5A**), allowing for direct comparison of the *RIF1* KO in ESCs to the *MCM10* mutants in iPSCs. *RIF1* ESC KOs were consistent among themselves (r = 0.97) yet significantly differed from WT controls (r = 0.56) (**Fig 5A**), showing variation across 56.6% of the genome in grouped analysis (39.5% and 37.6% individually), similar to HAP1 *RIF1* KOs. In contrast to a previous report (30), the replication profiles in ESC *RIF1* KOs (as well as HAP1 *RIF1* KO) did not appear random but instead were consistent among samples and differed from WT at well-defined sites (**Fig S14, Fig S17A**). Importantly, *MCM10* and *RIF1* showed different alterations in DNA replication timing (**Fig 5A**). When variant regions in these two gene mutations were merged, 94.2% of the genome demonstrated variant replication timing (**Fig S18**). Thus, the *MCM10* iPSCs we analyzed appear to harbor a previously undescribed alteration of the DNA replication timing program. These cell lines represent experimental repetitions and different iPSC clones derived from a single patient carrying compound heterozygous mutations in *MCM10* and characterized by natural killer cell deficiency (42). These *MCM10* mutations were previously shown to prevent its nuclear localization, causing de-stabilization of the replisome, reduced origin firing, genome instability and reduced cell proliferation (42, 43). Taken together, we propose that *MCM10* is a strong candidate for being a novel regulator of DNA replication timing.

**Fig 5.**
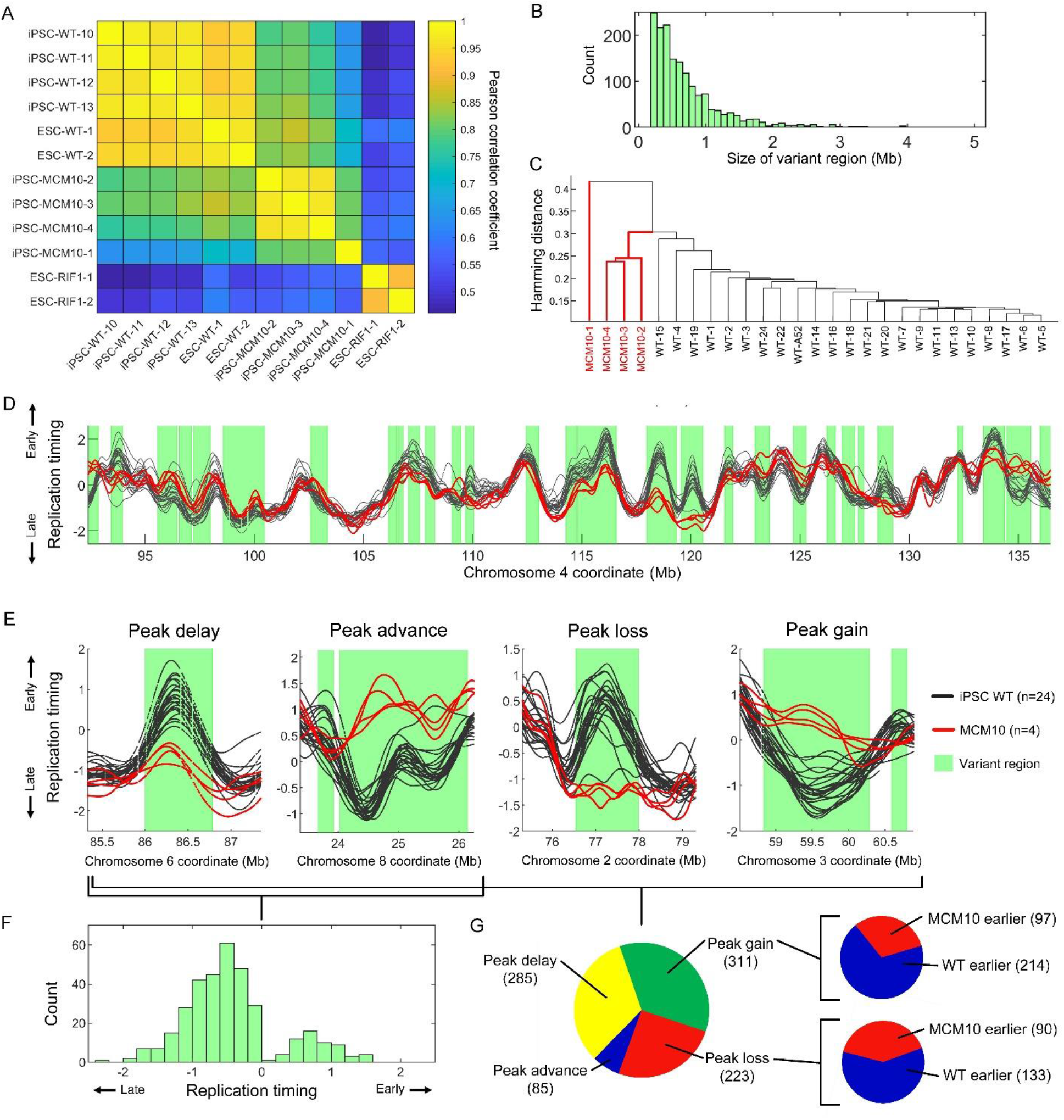
Mutations in *MCM10* are associated with extensive replication timing variation. (A) Whole genome correlation of the replication timing profiles of several WT iPSCs, *MCM10* mutants, ESC *RIF1* KO mutants, and normal ESC samples. (B) The distribution of the sizes of all *MCM10* variant regions. (C) Clustering of *MCM10*-1 and WT samples by peak presence. Sample *MCM10*-1 is an outlier, as it was in its genome-wide correlation values (**Fig S14**), indicating lower data quality; samples *MCM10*-3 and *MCM10*-4 are repetitions of the *MCM10*-1 cell line. (D) Example of replication timing variants in *MCM10*. (E) Examples of peak alterations in *MCM10* mutants. (F) The distribution of the change in replication timing at peaks within variant regions in *MCM10* relative to WT. (G) The number of peak delays, advances, losses and gains in *MCM10* mutants compared to WT, at *MCM10*-variant regions. Insets indicate the relative changes in replication timing at peak gain and loss sites.

*MCM10* replication profiles differed from the WT iPSC profiles across 1,613 variant regions, spanning 46.0% of the genome (13.4% to 45.8% in individual samples; **Fig S17B, Fig S19**). Variant regions spanned between 196Kb (a single sliding window in ANOVA) to a maximum of 4.9Mb, with a median of 532Kb (mean 713Kb; **Fig 5B**). Interestingly, we noticed that much of the variation in *MCM10* mutants localized to replication timing peaks – proxies for replication initiation sites (**Fig 5D, Fig S19**). Indeed, *MCM10* samples clustered separately from WT samples at peak locations (**Fig 5C**), indicating that a fundamental difference in replication between *MCM10* and WT cells resides at replication initiation sites.

To better understand how *MCM10* mutations influence replication initiation, we characterized four categories of peak change within *MCM10* variant regions: peak advance or delay, and peak gain or loss relative to WT (**Fig 5E**). Of 627 peaks shared between *MCM10* and WT, 285 showed replication timing delay while only 85 were advanced. This demonstrated substantial DNA replication initiation defects in *MCM10* mutant cells (**Fig 5C, F-G**). The median absolute change in replication timing in peak advances and delays was 0.65 units of standard deviation. Replication advances were more common in (but not exclusive to) later replicating parts of the genome, with the median peak advance having a WT (normal state) replication timing value of 0.42 units of standard deviation below the mean. Replication delays, on the other hand, were more common in very early replicating parts of the genome with the median delayed peak having a WT replication value of 1.29 units of standard deviation above the mean. Furthermore, *MCM10* demonstrated 311 peak gains and 223 peak losses relative to WT. At a majority of sites of either peak gain and loss, WT samples remained earlier replicating than *MCM10* (**Fig 5G**). Additionally, both peak gains and losses occurred more frequently at earlier replicating parts of the genome (**Fig S20**). The median peak gain site had a WT replication timing value of 0.77 and an *MCM10* replication value of 0.44 units of standard deviation above the mean. For the median peak loss, the replication timing values were 0.26 and 0.47, respectively.

In conclusion, impairment of *MCM10* appears to exert a global influence on genomic replication timing, in particular perturbing normal replication initiation. This included both replication delays at sites of shared initiation between WT and *MCM10* cells; as well as gains and losses of replication initiation sites, predominantly at early replicating genomic regions.

## Discussion

Identifying genetic alterations that lead to reprogramming of DNA replication timing can illuminate the molecular mechanisms of DNA replication control. However, almost no such factors have been identified to date despite intensive efforts. Here, we took, to our knowledge, the most comprehensive characterization of replication timing alterations in gene knockouts and genetic diseases. Apart from the previously described role of *RIF1* in replication timing control, we identified a novel role for *MCM10* and a possible albeit potentially indirect involvement of the *BLM* helicase in DNA replication timing. *MCM10* is a conserved and essential component of the DNA replication initiation machinery (44) and we show here that disease-associated mutations in the *MCM10* gene lead to extensive perturbation of origin firing genome-wide.

Notably, we did not observe similar replication timing aberrations in cells mutated for other central components of the DNA replication initiation machinery, such as *GINS4* and *RECQL4*. The replication timing phenotype of *MCM10* mutant cells therefore appears to be highly specific, consistent with the diverse roles of *MCM10* in stabilizing the CDC45:MCM2-7:GINS helicase as well as the proceeding replisome (43). Further research will be required to better understand the mechanisms by which *MCM10* may control replication initiation timing as well as to link *MCM10* mutations to genome stability and cellular and disease phenotypes. In particular, although we identify a consistent replication timing phenotype across different experimental repetitions and patient-derived iPSC mutant clones of *MCM10*, they are all derived from the same individual. Identification of additional individuals with *MCM10* mutations and complementary studies using engineered cell lines (43), will further establish the role of *MCM10* in replication timing. However, naturally occurring mutations in *MCM10*, such as the compound heterozygote analyzed in this study, are extraordinarily rare.

Although we ultimately ruled out most tested genes, the majority of tests initially resulted in the positive identification of changes in DNA replication timing. We show that this is expected given the genome-wide nature of variant search, which is especially pronounced in the case of replication timing data due to its chromosomal continuity. We thus emphasize the need for rigorous consideration of multiple testing in any genome-wide search for replication timing alterations in any biological system. In contrast, many previous studies used arbitrary thresholds for identifying variants and determining whether a gene mutation influences DNA replication timing. By applying an empirical false-discovery correction, we were able to rule out many candidate replication timing regulators, including some that were proposed by previous studies.

A main limitation of our study is that we focused on three specific cell types, among which LCLs and HAP1 cells are both transformed or derived from tumor cells, while iPSCs are pluripotent stem cells. It is conceivable that alterations in replication timing would only be observed in certain normal cell lineages. This is consistent with the specific symptoms and affected tissues in patients with replication gene mutations. Thus, it remains of interest to study replication timing alterations in different genetic backgrounds in a variety of differentiated cell types. Furthermore, the effects of human disease-associated variants may differ depending on the relative pathogenicity of the variant. Another limitation of our study is the use of bulk cell samples for analysis. Newer single cell approaches (45) have the potential to reveal stochastic events that are specific to individual cells rather than being shared across a population of cells. For instance, a genetic mutation may affect the activity of different replication origins or replication forks in each cell, thus evading detection in bulk analysis yet still impacting tissue physiology in an affected human. Notwithstanding these caveats, we find it remarkable that the replication timing program is so robust to a wide array of genetic perturbation in *trans*. While *cis*-acting polymorphisms can affect local replication timing in different cell types and across many genomic loci (19, 20), it appears that global changes in replication timing may not be compatible with cell or human health. It is intriguing to consider the possible reasons for this rigidness of the replication timing program. In particular, interactions of replication timing with gene regulation and with genome stability – and even the intersection of the two (46) – may define DNA replication timing as an essential cellular program.

## Methods

### Cell lines

#### LCL

We analyzed 271 LCL samples from 245 individuals. These samples covered 32 genetic diseases across 117 individuals (134 total samples) and 124 individuals (137 samples) of presumed healthy status (**Table 1**). 138 of the LCL samples were obtained from the Coriell Institute for Medical Research (Camden, NJ) as either DNA samples or cell cultures. Five FA mutant cell lines were obtained from the International Fanconi Anemia Tissue and Cell Repository housed at the Rockefeller University. LCLs were cultured in Roswell Park Memorial Institute 1640 medium (Corning Life Sciences, Tewksbury, MA, USA), supplemented with 15% fetal bovine serum (FBS; Corning). Culture was maintained at 37°C with 5% CO2 in a humidified atmosphere. Sample identification numbers and genotypes are available in **Supplemental Table 1**.

Among the remaining samples, the repeat expansion cohort provided 55 mutant LCL samples from six diseases and 48 presumed healthy samples (47). The Illumina platinum family provided 17 presumed healthy samples (48).

#### HAP1

HAP1 WT and KO cell lines were obtained from Horizon. The cell lines were cultured following the provider’s recommendations, in IMDM medium supplemented with 10% FBS and 1% Penicillin/Streptomycin. Culture was maintained at 37°C with 5% CO2 in a humidified atmosphere. Cells were passaged every 2-3 days.

Cells were harvested at 70-80% confluence with 0.05% trypsin at 37°C for 5 min. Dissociation of cells was checked using microscopy. Cells were split into two samples containing approximately 2×10^6^ cells each, one of which was used for FACS analysis and the other for DNA extraction. Cells collected for FACS analysis were pelleted at 1000 rpm at 4°C for 5 min, washed once in 500 ml of ice-cold PBS and resuspended in 250 μL of ice-cold PBS. Cells were fixed in 750 μL of ice-cold (−20°C) ethanol with constant gentle vortexing and stored at 4°C. Fixed cells were washed with 400 μL of ice-cold PBS and centrifuged at 1000 rpm at 4°C for 5 min. Cells were resuspended in 400 μL of PBS, 400 μL of Accutase and incubated at room temperature for 20 min. Cells were then pelleted and resuspended in 400 μL of PBS. RNase A (10 mg/ml) treatment was done at 37°C for 30 min. Propidium iodide at a concentration of 1 mg/mL was added and staining was done at room temperature in the dark for 30 min. Cells were passed through a polystyrene cell-strainer-capped tube before flow cytometry analysis. Analysis of PI-stained cells was done with a FACSAria Fusion Sorter. Cells of known ploidy (haploid/diploid) were used as controls for each run. All HAP1 KO samples were sequenced as either diploidized samples, or both as haploid and diploidized samples. In those cases where both haploid and diploid samples were analyzed from the same KO clone, no significant differences were observed in replication timing as a function of ploidy, and the two samples were considered experimental repetitions of the same KO clone.

#### iPSC

The iPSCs from *MCM10-, CDC45*-, and the *GINS4*-deficient individuals were cultured in feeder-free condition with Stemflex (Gibco, A3349401). iPSC lines for RTT, RTTC, LNS and matched WT controls were obtained from the Coriell Institute (see **Supplemental Table 1**). iPSC lines GM27622 and GM27629 were grown in mTeSR Plus Basal medium and iPSCs for FXS and matched controls (49, 50) were grown in mTeSR1 medium (STEMCELL Technologies). Cell lines GM27437, GM26077, GM27730 and GM260105 were grown on a layer of Mouse Embryonic Fibroblast feeder cells (Gibco) coated with D-MEM/F-12 supplemented with 20% of KnockOut Serum replacement (Gibco) and 10 μg/mL of Basic Fibroblast Growth Factor (bFGF) (Gibco, PHG0264). Feeder-dependent cells were transferred and adapted to Matrigel conditions following the recommendations of STEMCELL Technologies. All cell lines were grown at 37°C, 5% CO2, and passaged by dissociating to single cells with Accutase (Sigma, A6964) and plating at a density of 1×10^6^ cells/well in Matrigel-coated 6 well plates. For the first 24 hours after passaging, cells were cultured with 10μM ROCK inhibitor (Y-27632; STEMCELL Technologies).

#### ESC

The WT and *RIF1* KO human stem cells were cultured in StemFlex media (Thermo Fisher A3349401) on Geltrex (Thermo Fisher A1413302). Upon reaching 70% confluency, cultures were passaged at a ratio of 1:10, or cryopreserved in a solution of freezing media containing 40% FBS (Gemini Bio-Products 900-108) and 10% DMSO (Sigma Aldrich D2650). Passaging was performed by TrypLE (Life Technologies 12605036) dissociation to small clusters of cells, and plated in media containing 10 μM Rock inhibitor Y-27632 (Selleckchem S1049) was added to media and removed within 24-48 h. Cells beyond passage 10 were no longer supplemented with Rock inhibitor. All embryo and ESC research was reviewed and approved by the Columbia University Embryonic Stem Cell Committee and the Institutional Review Board.

In preparation for CRISPR, guide RNAs were designed targeting relevant genes using software from cripsr.mit.edu or Benchling.com. Guides were chosen with the highest index score in a region closest to the DNA region of interest. Nucleofection was performed using Amaxa Cell Line Nucleofector Kit II, program A-23 with a Cas9-GFP plasmid (Addgene, 44719) and guide RNA. Cells were plated, cultured for 2 additional days, stained with 10 μg/mL Hoechst 33342, and subsequently sorted via FACS for cells that had both haploid DNA content and GFP positivity. Single colonies were propagated, and duplicates were made for cryopreservation and for DNA isolated for PCR and Sanger sequencing.

Fluorescence-activated cell sorting (FACS) was performed using the FACS-Aria machine at the Columbia University Stem Cell Initiative flow cytometry core. Populations were gated first for cells, followed by gating for single cells. Cells were suspended in media containing 10% FBS in PBS (Life Technologies 14190-250) throughout the analysis. Live cells were kept on ice during transportation and analysis. Analysis was performed using FloJo software (BD Biosciences).

### Generation of replication timing profiles

DNA was extracted using the MasterPure™ Complete DNA and RNA Purification Kit (Lucigen) following the manufacturer’s instructions. PCR-free whole genome sequencing was performed using paired-end reads (GeneWiz, South Plainfield, NJ, see **Supplemental Table 1**). Sequencing reads were converted into non-mapped bam files and marked for Illumina adaptors and duplicate reads with Picard Tools (v1.138) (http://broadinstitute.github.io/picard/) commands ‘FastqToSam’, ‘MarkIlluminaAdapters’, and ‘MarkDuplicates’. Bam files were aligned to hg19 with BWA mem (51) (Li 2013) (v0.7.17). GC-corrected read depth data for each sample were then generated via TIGER (36) using a read length of 36bp for alignability filtering and a bin size of 2500bp. All other parameters were TIGER defaults.

The raw post-GC-corrected data was then filtered for copy number alterations using permissive parameters such as to retain replication timing information and any potential disease-related replication timing alterations. To remove clonal or sub-clonal aneuploidies, individual autosomes were first removed if they had a copy number > 2.2 or < 1.8. This step removed whole chromosomes from diseased samples with known aneuploidies (e.g., TRI21) along with healthy samples with sub-clonal aneuploidies. Notably, chromosome 1q is removed in analyses of all *MCM10* and HAP1 due to high copy number (all *MCM10* mutants and 50% of HAP1 WT samples were affected). Next, regions of large (>1Mb) duplications or deletion were manually removed by visually comparing the distribution of raw data across all samples. To filter outlier and smaller CNVs, windows 4 standard deviations above or below the mean copy number per chromosome were removed. Each sample was then filtered via the TIGER command ‘TIGER_segment_filt’ (using the MATLAB function ‘segment’, R2 = 0.04, standard deviation threshold = 2.5). These steps optimally corrected samples across all cell types and data qualities.

Replication timing values were generated by smoothing the filtered GC-corrected data with a cubic smoothing spline (MATLAB command ‘csaps’, smoothing parameter = 1×10^-17^). Only regions of >20 continuous 2500bp windows were included and smoothing was not performed over data gaps >100kbp or reference genome gaps >50kbp. The smoothed profiles were then normalized to an autosomal mean of zero and a standard deviation of one.

### Detection of replication timing variant regions via ANOVA

Analysis of variance (ANOVA) was performed on autosomes to detect regions of replication timing variation among samples. We performed the variant analysis both for individual mutant samples against all WT samples of the same cell type and again with samples grouped by mutated gene. For the *RIF1* KO of ESCs, WT iPSCs were used. To avoid regions with different numbers of analyzed samples due to sample-specific filtered regions or chromosomes, we substituted individual windows of WT samples where less than 5 samples have missing data with the average filtered unsmoothed data of the WT samples. All filtered unsmoothed data was then mean-shifted to an autosomal genome copy number of 2.

We performed one-way ANOVA in a sliding window of 76 x 2500bp bins (185kb window) with a quarter step of 19 bins (47.5kb) on the filtered GC-corrected unsmoothed data (40). The corresponding p-value for each window was calculated with the MATLAB function ‘anova1’. ANOVA was not performed in windows with complete missing data for one or more mutant samples to avoid a local sample number disparity. We called variant regions as windows with a p-value less than the Bonferroni-corrected threshold based on the number of ANOVA tests performed for each individual mutant sample or mutated gene group. Adjacent significant windows were merged, and the p-value was recalculated over the merged region (in later analyses for FDR calculation, only the non-merged windows were used). The proportion of the genome with variant replication timing was then calculated from the total length of regions assigned as variant divided by the length of the genome analyzed.

### Comparison of replication timing profiles, PC analysis, and age analysis

All replication timing profile correlation was calculated as Pearson’s correlation coefficients. In comparing the TIGER and S/G1 replication timing profiles of sample NA12878, S/G1 coordinates were lifted from hg38 to hg19 with vcf-liftover (https://github.com/hmgu-itg/VCF-liftover) and interpolated to TIGER window coordinates with the MATLAB function ‘interp1’.

PC analysis was performed on the replication timing profiles of all autosomes with the MATLAB function ‘pca’. To determine if age influenced DNA replication timing, PC analysis was only performed on 83 WT individuals with known age. In calculating the relationship between age and replication timing correlation, the median correlation of a sample to all other samples was used. A linear model was fit using the MATLAB function ‘fitlm’. The linear model excluded samples with a correlation ≤ 0.7 to all other LCL samples. However, when including all samples in the analysis, the linear correlation was still insignificant (**Fig S3B**). Although including all samples produced a marginally higher correlation (r^2^ = 2.75 x 10^-3^ to r^2^ = 0.189), this was driven by a few samples from young individuals with abnormal replication profiles.

In comparing repli-seq profiles of ESC and differentiated cells to *MCM10*, coordinates were interpolated to TIGER window coordinates with the MATLAB function ‘interp1’.

### P-value inflation analysis and ANOVA simulations

For each mutant-WT ANOVA test, the slope of the QQ-plot relationship of theoretical vs sample quantiles was extracted with the MATLAB function ‘fitlm.’ To determine FDR for each analysis, q-values were calculated by the MATLAB function ‘mafdr’ using the Benjamini-Hochberg method (52). FDR for each ANOVA test was determined as the proportion of q-values less than the original Bonferroni-corrected p-value threshold compared to its original p-value (i.e., the proportion of windows identified as false positive) (**FigS4**).

ANOVA simulations used all samples with a >0.7 correlation to all other samples regardless of mutant or WT status. The simulations were performed for different combinations of sampled mutant and WT samples. In 1000 iterations for each combination, samples were randomly assigned into the WT or mutant groups and an ANOVA test was performed identically to the true mutant-WT tests. For each iteration, the proportion of the genome with variant replication timing and the p-values for each window were analyzed.

### Overlap in regions of replication timing variability

Overlap percentage was calculated in pairs of all individual mutant samples (184 x 184 tests) where the variant regions of one query sample were compared to the variant regions in one subject sample. For each analysis, the number of overlapping nucleotides in the variant regions of the query and subject sample were calculated with BEDtools intersect (53). The overlapping percentage was determined as the number of overlapping nucleotides divided by the total length of variant replication timing regions for the query sample. Therefore, the overlap percentage can differ for two samples depending on which is the query or subject sample. For the significance of overlap, a Fisher’s exact test was performed on pairs of individual mutant variant regions using BEDtools fisher. The p-value from the two-tailed t-test was used.

For determining the proportion of the genome with variant replication timing covered by both *MCM10* and *RIF1* KO in ESCs, variant regions were merged with BEDtools merge. The sum of the merged variant regions was divided by the length of the genome available for analysis in *MCM10* (which was shorter than *RIF1* by 7.12 Mb).

### Identifying and clustering replication timing peaks

Chromosome 1p was removed for all peak analyses as it was filtered out in *MCM10.* For identifying peaks in *MCM10* and iPSC WT samples, the pairwise distances of local maxima in the individual mutants were calculated with the MATLAB function ‘pdist’. Hierarchical clustering was then performed on the pairwise distance matrix using the MATLAB function ‘linkage’ using the average method and the default metric of Euclidean distance. Peak clusters and ranges were next calculated with the MATLAB function ‘cluster’ using a cutoff of 20,000bp as the distance criterion for forming clusters. For determining peak overlap, only peaks present in at least 75% of *MCM10* or WT samples were compared. A peak was considered shared if the range of an *MCM10* peak and WT peak overlapped as calculated by BEDtools intersect.

In determining peak advances or delays, only peaks overlapping *MCM10* variant regions were considered. The relative change in replication timing was determined as the change in mean replication timing of *MCM10* samples to WT samples within the shared peak range. For peak gains or losses, only peaks present in at least 75% of *MCM10* or WT samples were compared. Peak gains were defined as peaks present in *MCM10* but not WT and peak losses were defined as peaks present in WT but not *MCM10*. Replication timing changes in peak gains and losses were calculated within the range of the peak cluster using either the mean WT value or mean *MCM10* value, as applicable.

For clustering *MCM10* and WT samples by peak use, all 6234 peaks present in any of the samples were included. The binary presence or absence of the 6234 peaks in *MCM10* and WT samples was clustered with the MATLAB function ‘linkage’ using the average method and the metric of Hamming distance (the percentage of coordinates that differ).

## Data availability

Raw sequence data is available on SRA under the bioproject PRJNA754107 (HAP1 samples and iPSC and LCL samples approved for non-restricted data access) and on dbGaP with accession numbers phs001957 (ESCs) and phs002597 (iPSCs and LCLs).

## Acknowledgements

This work was supported by NIH grants 1DP2GM123495 (to A.K.), R01GM123018 (to M.B.S.), NYSTEM IDEA award #C32564GG (to D.E.), and in part by NIH-NIAID R01AI137275 (to E.M.M). MCM10 and GINS4 iPSCs were a gift from Dr. Jordan Orange and generated through the support of NIH-NIAID R01AI120989. Fanconi Anemia cell lines were obtained from the International Fanconi Anemia Tissue and Cell Repository housed at the Rockefeller University.

## Notes

### Competing Interest Statement

The authors have declared no competing interest.

### Summary of Updates

Funding source added

