## Supplemental Figures for "Comprehensive analysis of DNA replication timing in genetic diseases and gene knockouts identifies *MCM10* as a novel regulator of the replication program"

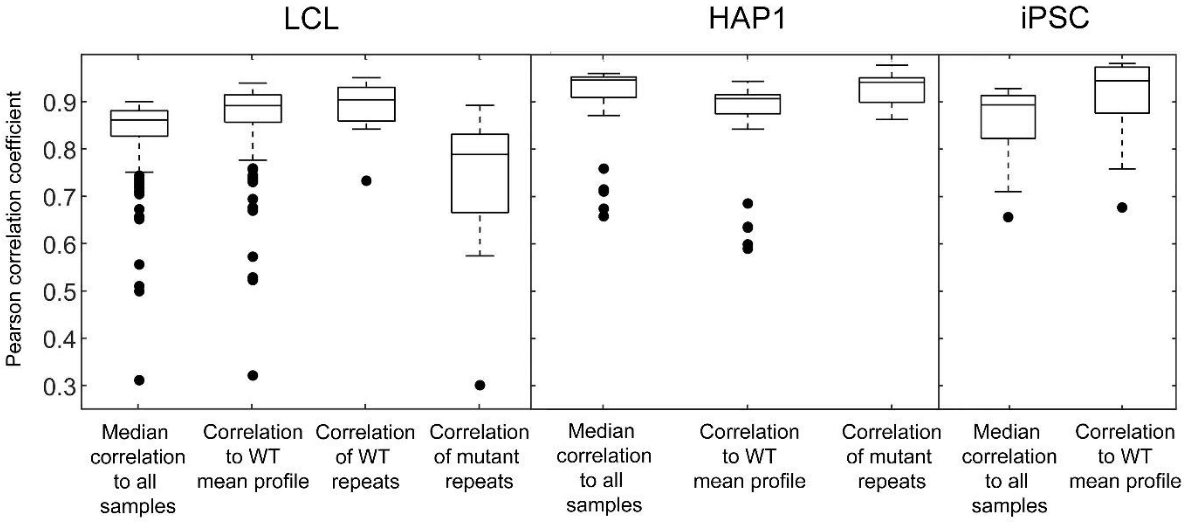


**Fig S1. Correlation of replication timing profiles.** Different correlation-based metrics of samples: median correlation of all samples of the same cell type, correlation of samples to the mean WT profile of the respective cell type, and correlation of repeat samples (genetically identical samples in LCL and samples with the same gene KO in HAP1).

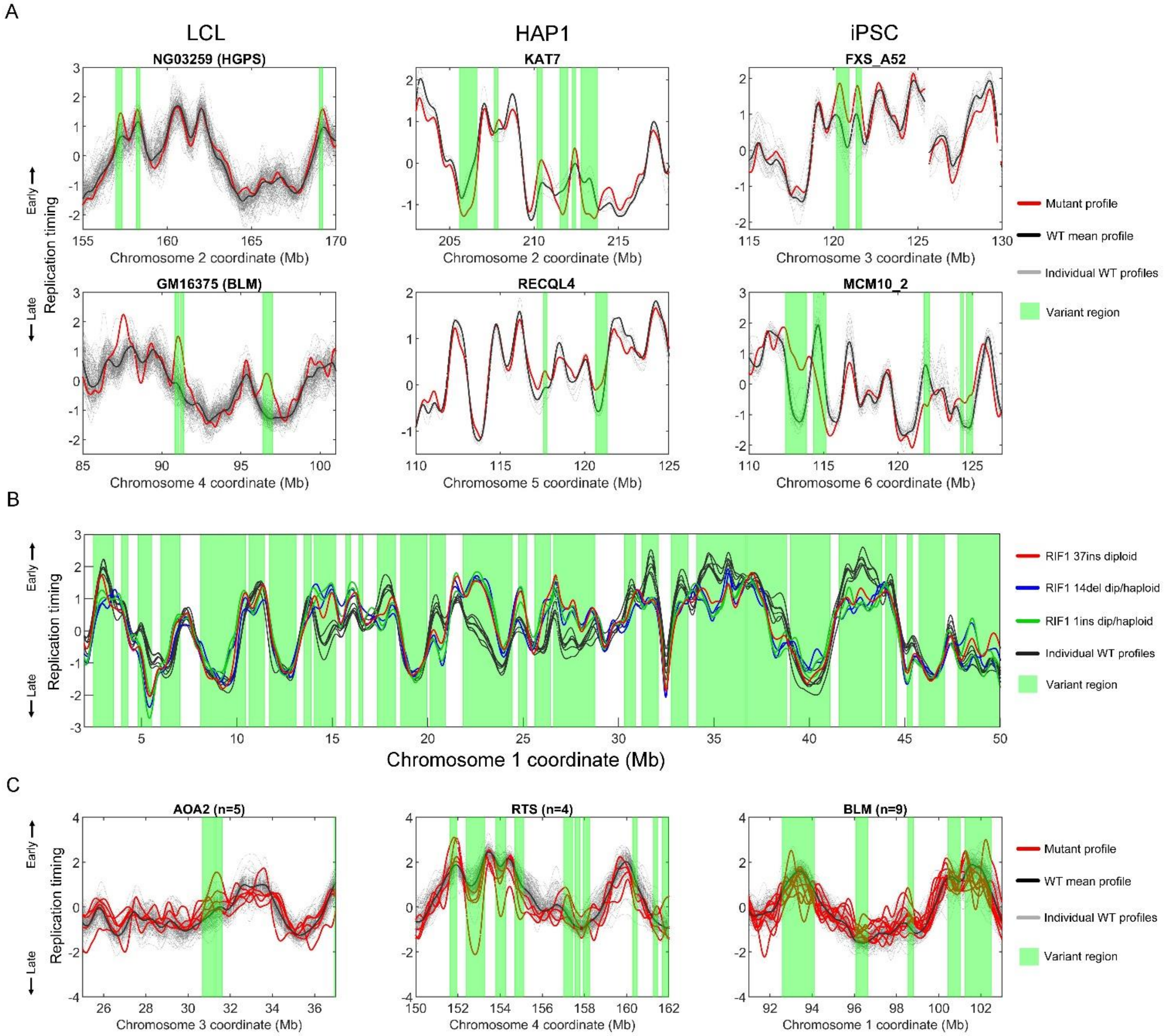


**Fig S2. Additional examples of mutant sample replication timing.** (A) Examples of variant replication timing in individual samples. (B) HAP1 KO of RIF1 compared to WT HAP1 replication timing. Samples are colored by RIF1 gene mutation with two mutant cell lines each having both a haploid and diploid sample. (C) Samples grouped by mutated gene where variant replication timing was driven by a small number of abnormal samples.

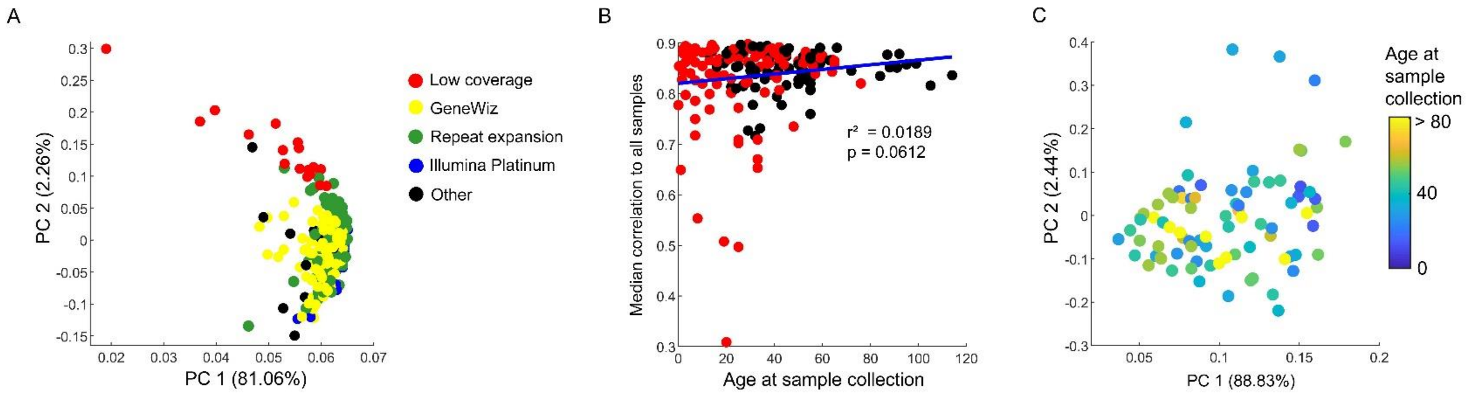


**Fig S3. Stratification of variant replication timing by sequencing cohort and age.** (A) PC analysis of replication profiles in LCL by sequencing cohort. (B) Relationship of individual age to median correlation of autosomal replication timing among all LCL samples. (C) PC analysis of replication timing for 83 WT LCL samples with known ages.

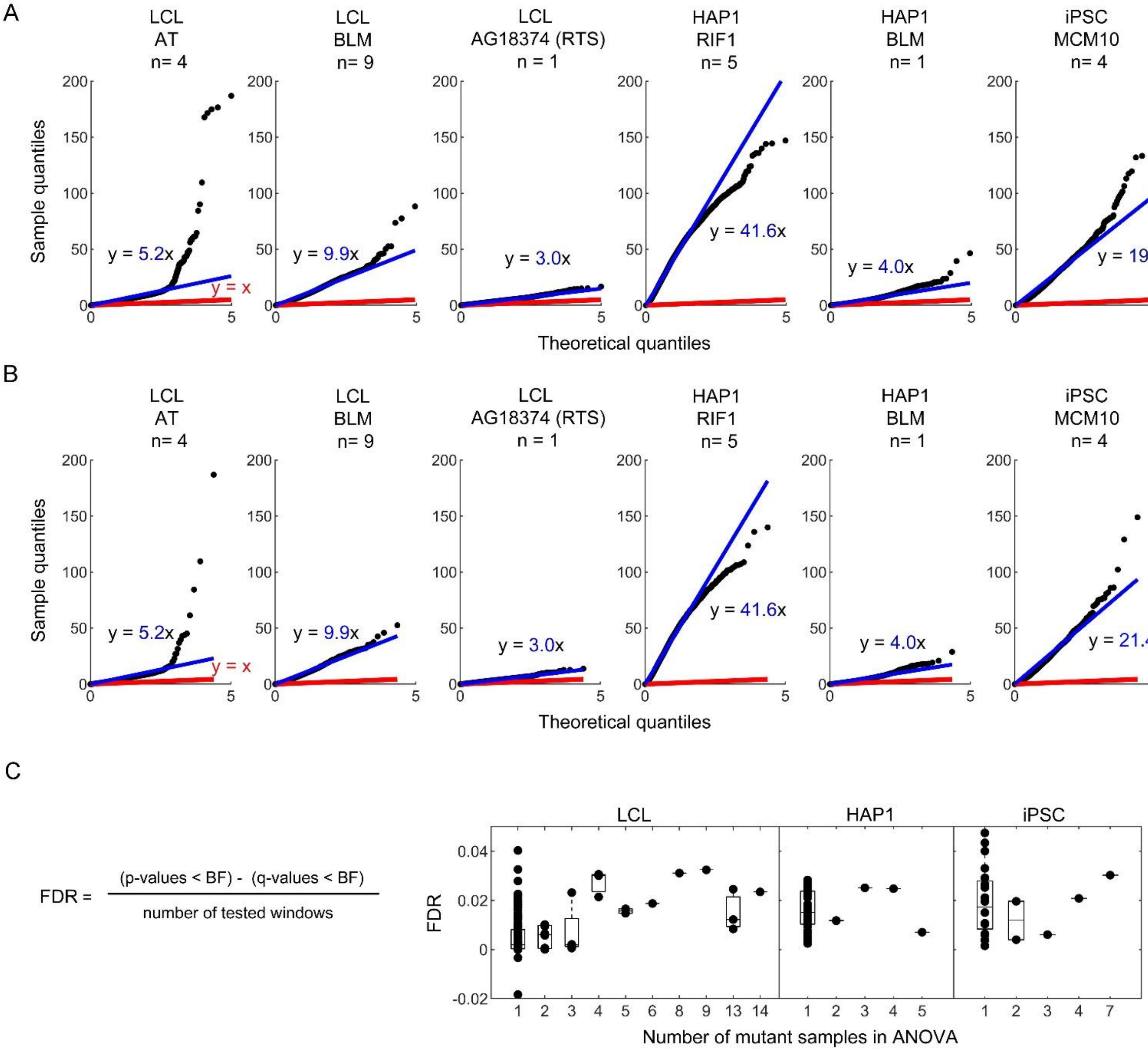


**Sig S4. P-value inflation of ANOVA.** (A) Additional QQ plots of ANOVA p-value inflation from samples grouped by mutated gene or as individuals. (B) Identical analysis as (A) but using non-sliding windows in ANOVA. P-value inflation is still observed in independent windows. (C) q-values are calculated using the Benjamini-Hochberg procedure in each ANOVA test. The proportion of reassigned q-values from the original p-values that are smaller than the Bonferroni-corrected (BF) threshold is the false-discovery rate (FDR) in each test.

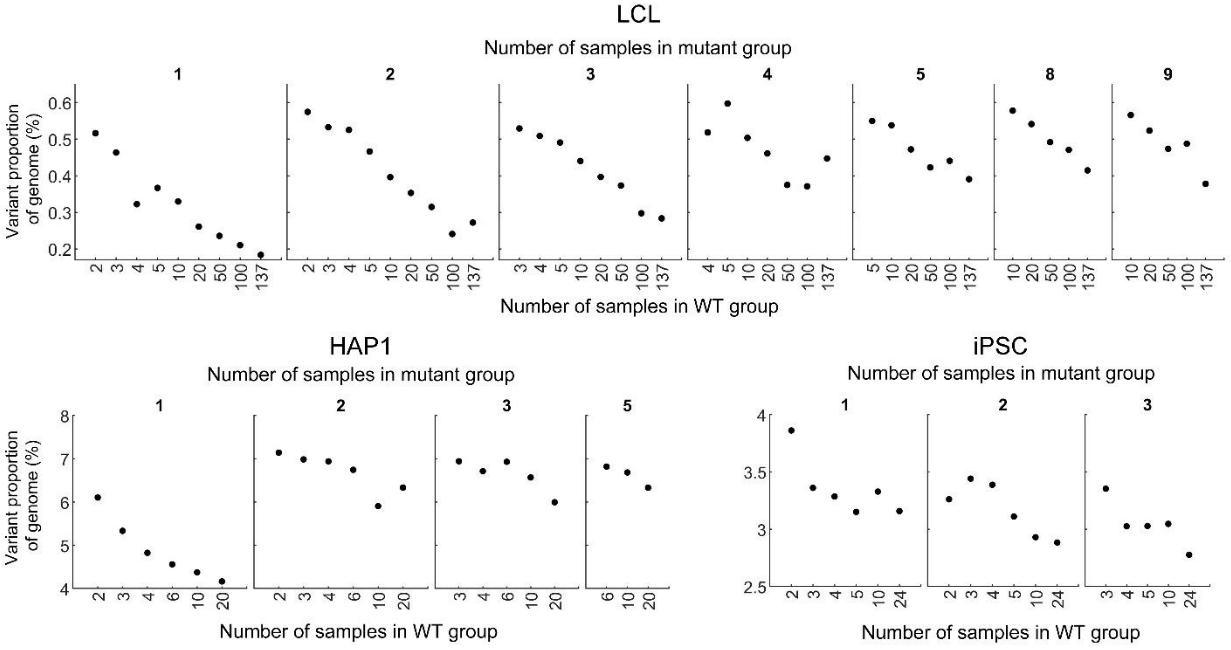


**Sig S5**. **Implementation of simulation false discovery rates.** Median proportion of the genome that is observed as variant for replication timing in 1000 ANOVA simulations of mutant and WT groups with different number of samples. See Fig S6 for the full distributions in all mutant group sample numbers.

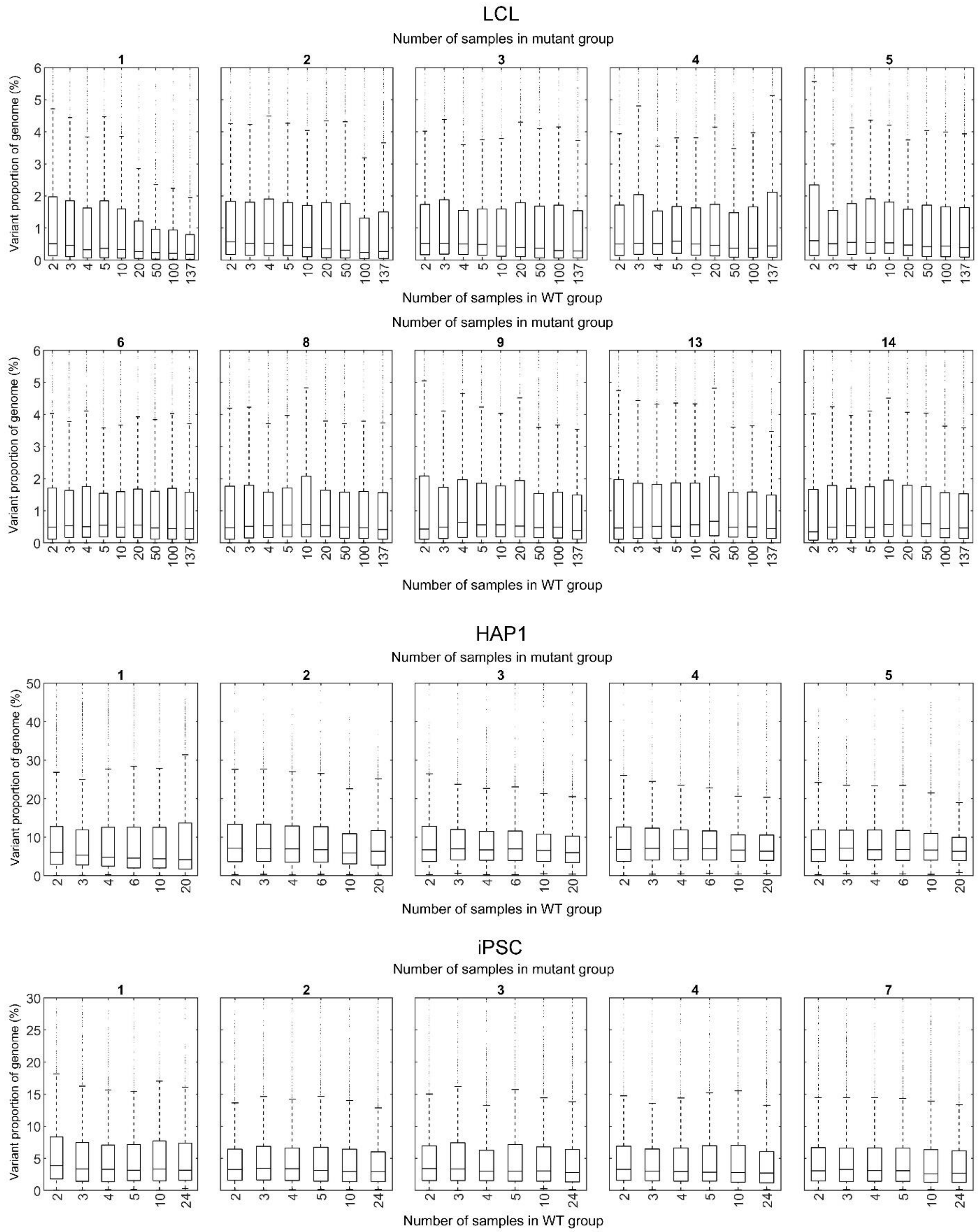


**Sig S6**. **Boxplot of simulation results.** Variant genome proportions from 1000 ANOVA simulations of different sized mutant and WT groups.

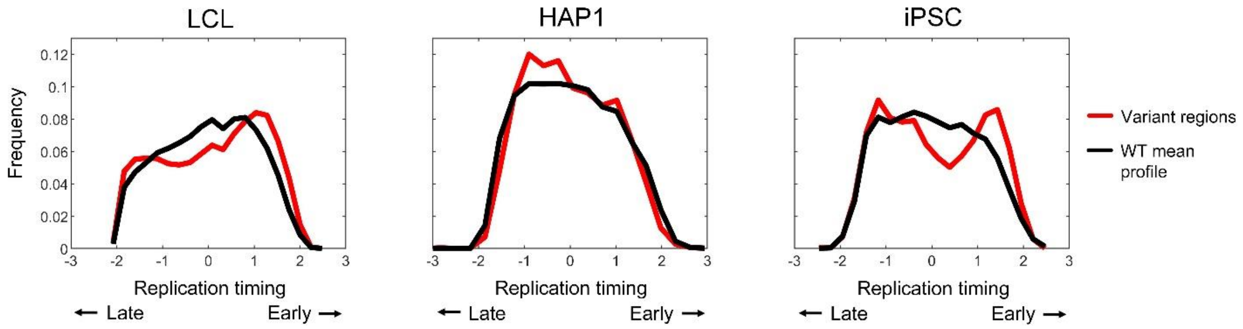


**Fig S7**. **Distribution of variant region replication timing in mutant analyses.** Histograms of the mean replication timing for variant regions in all mutated gene groups and individuals compared to the genome-wide distribution of replication timing values for the WT mean profile.

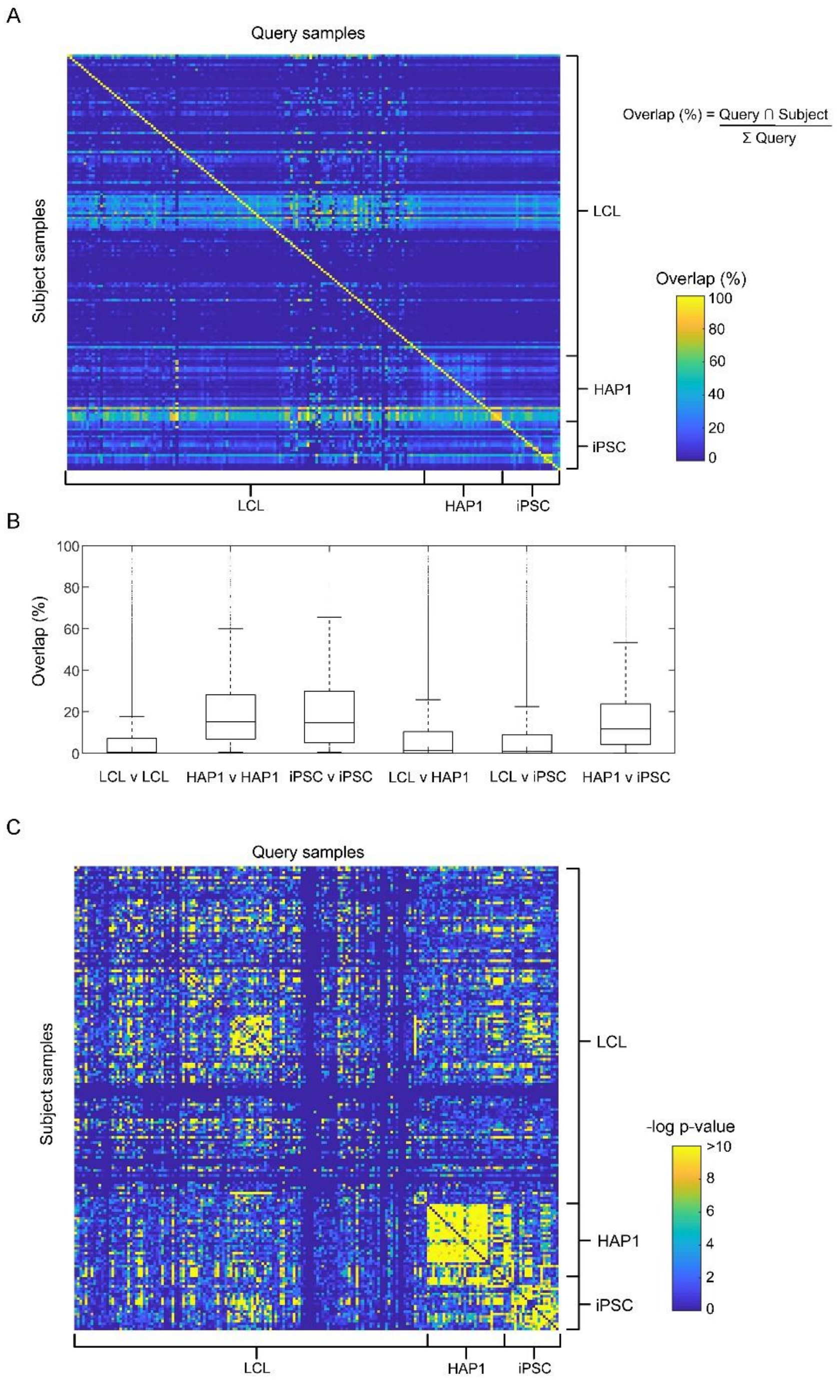


**Fig S8**. **Overlap of regions with replication timing variability in individual mutated samples**. (A) The extent of overlap (in percent) of variant regions for all individual mutant samples calculated as the proportion of nucleotides shared between any query and subject sample divided by the total length of variant regions in the query sample. All samples can be either the query or subject with the query serving as the reference for the overlap statistic. Regions of high overlap correspond to samples with high variant replication timing proportions. (B) Boxplot of the percent overlap within and between cell types from (A). (C) Significance of overlap of variant regions for all individual mutant samples calculated from a Fisher's exact test. Tests comparing the same samples (diagonal line) were removed.

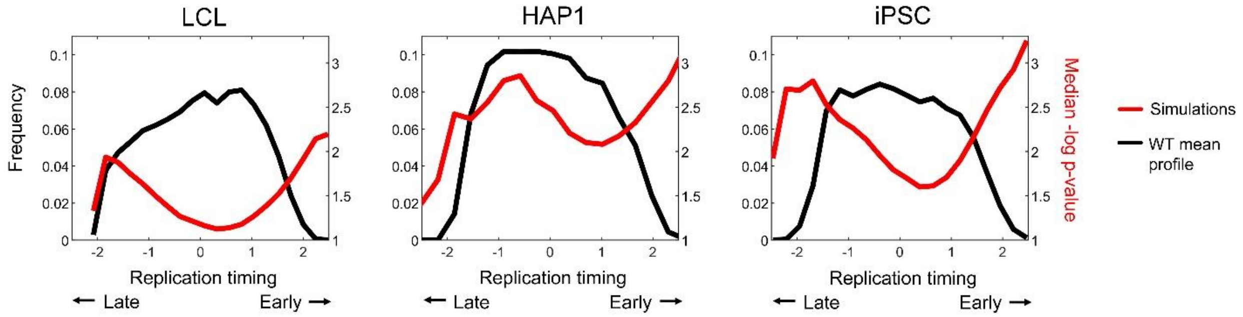


**Fig S9**. **Distribution of p-values in simulations of replication timing variation**. Histograms of replication timing values for the WT mean profile (black) overlayed with a histogram of median p-values from all simulations within the same bins used for replication timing (red).

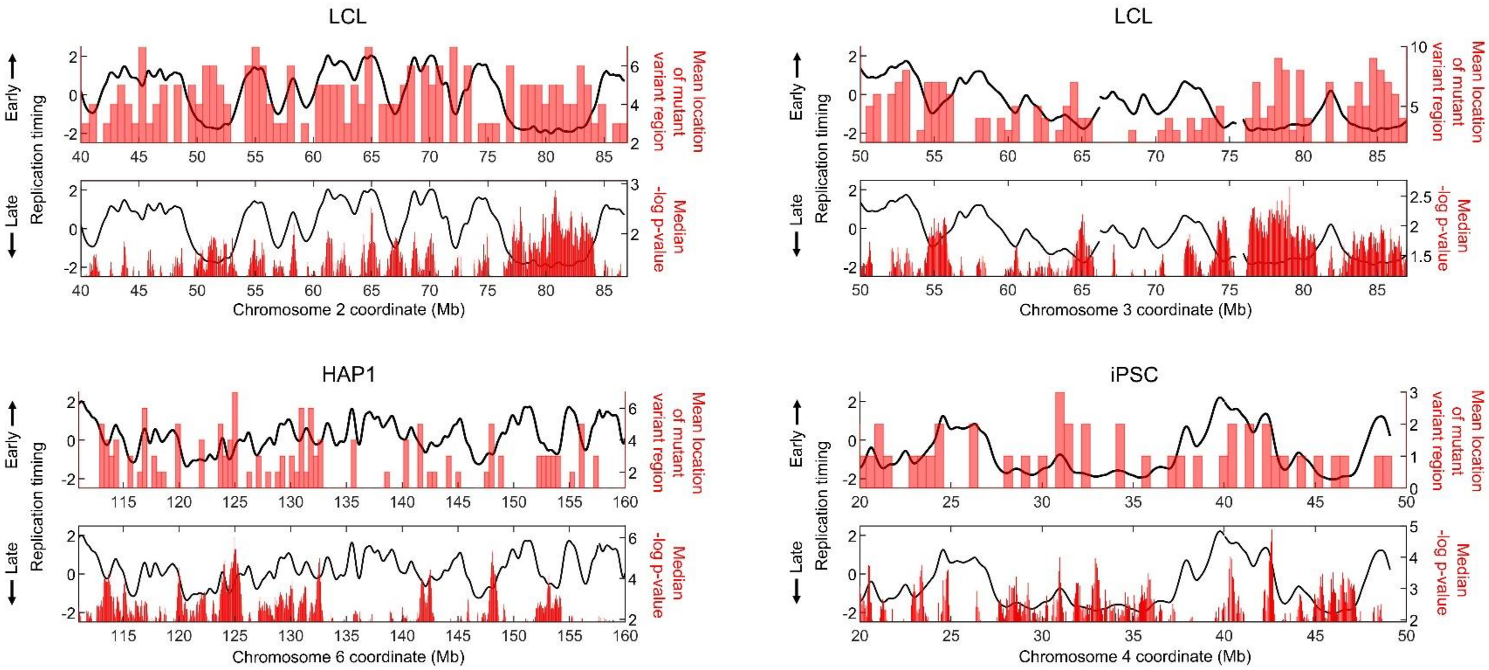


**Fig S10**. **Comparison of variant regions in simulations and actual mutant analyses.** Shown are two chromosomes for LCLs and one chromosome each for HAP1 and iPSCs. In each, the top plots show the locations of variant regions across all mutated gene group and individual analyses, while the bottom plots show the median p-value across 1000 simulations. All mutant group sizes were used, tested against the total number of WT samples. Only p-values below the median are shown.

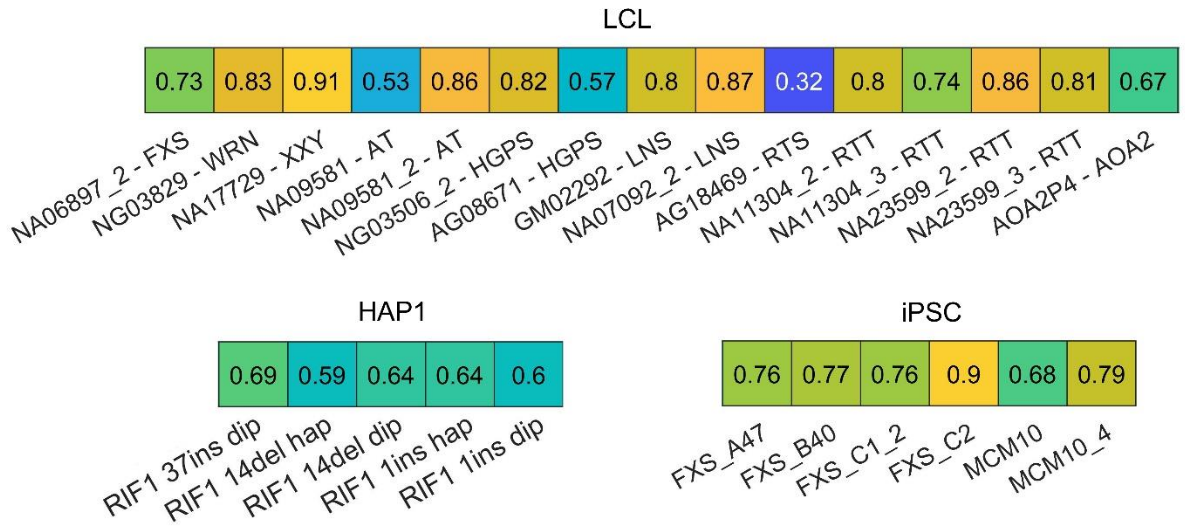


**Fig S11**. **Correlation of individual mutant samples to the mean WT profile**. The correlation of individual mutant samples with high replication timing variability to the mean WT replication timing profile in each respective cell type.

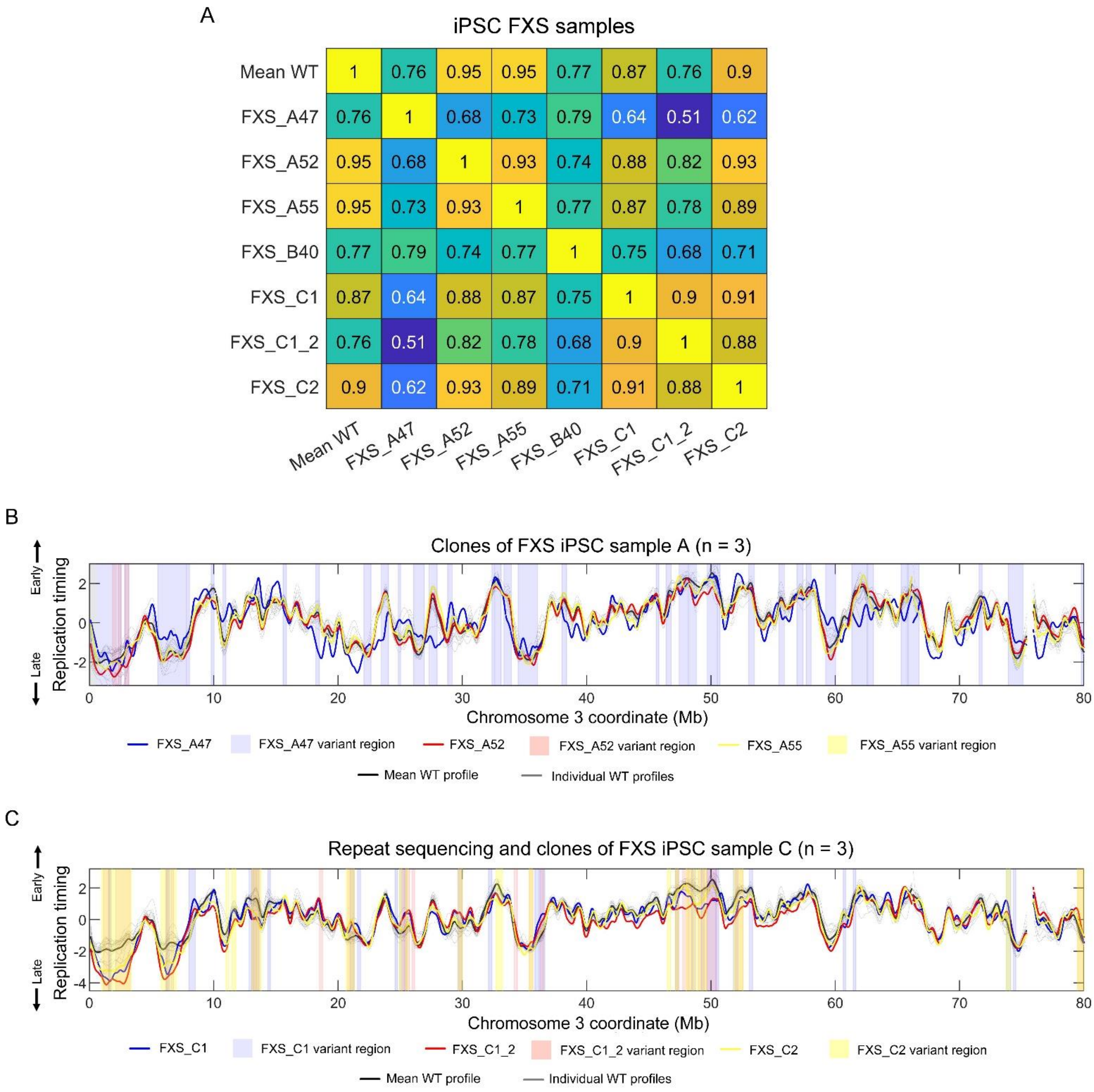


**Fig S12**. **Variant replication timing of individual FXS samples**. (A) Correlation of all iPSC FXS samples to the mean WT iPSC profile. (B) Replication timing profiles and variant regions for three independent clones of iPSC FXS sample A. FXS_A47, FXS_A52, and FXS_A55 are all clones of the same FXS donor. (C) Replication timing profiles and variant regions for two independent clones of iPSC FXS sample C. FXS_C1_2 is a re-sequencing of FXS_C1. FXS_C1 and FXS_C2 are clones of the same FXS donor.

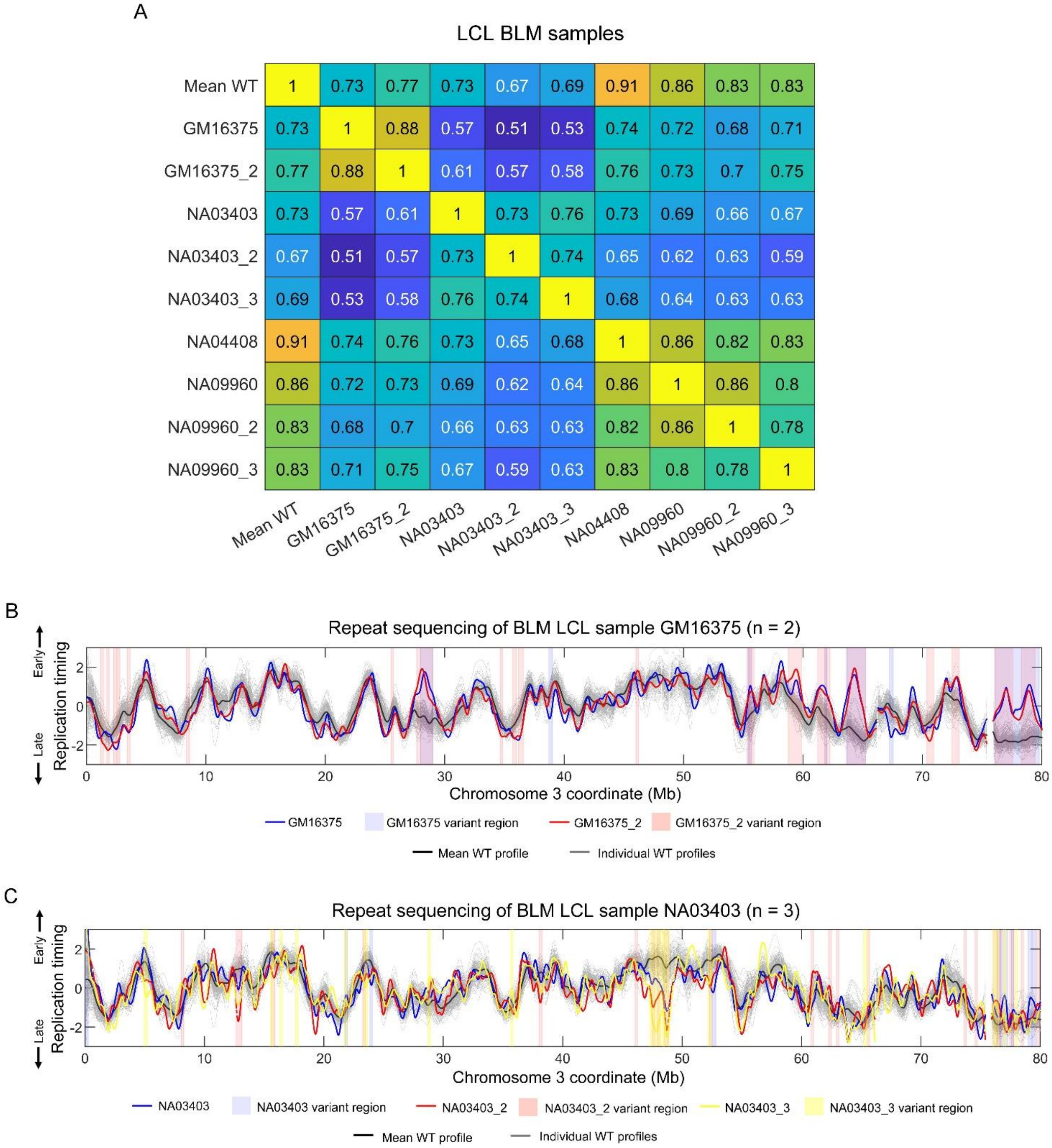


**Fig S13**. **Variant replication timing of individual BLM samples.** (A) Correlation of all LCL BLM samples to the mean WT LCL profile. (B) Replication timing profiles and variant regions for the resequenced BLM sample GM16375. (C) Replication timing profiles and variant regions for the resequenced BLM sample NA03403.

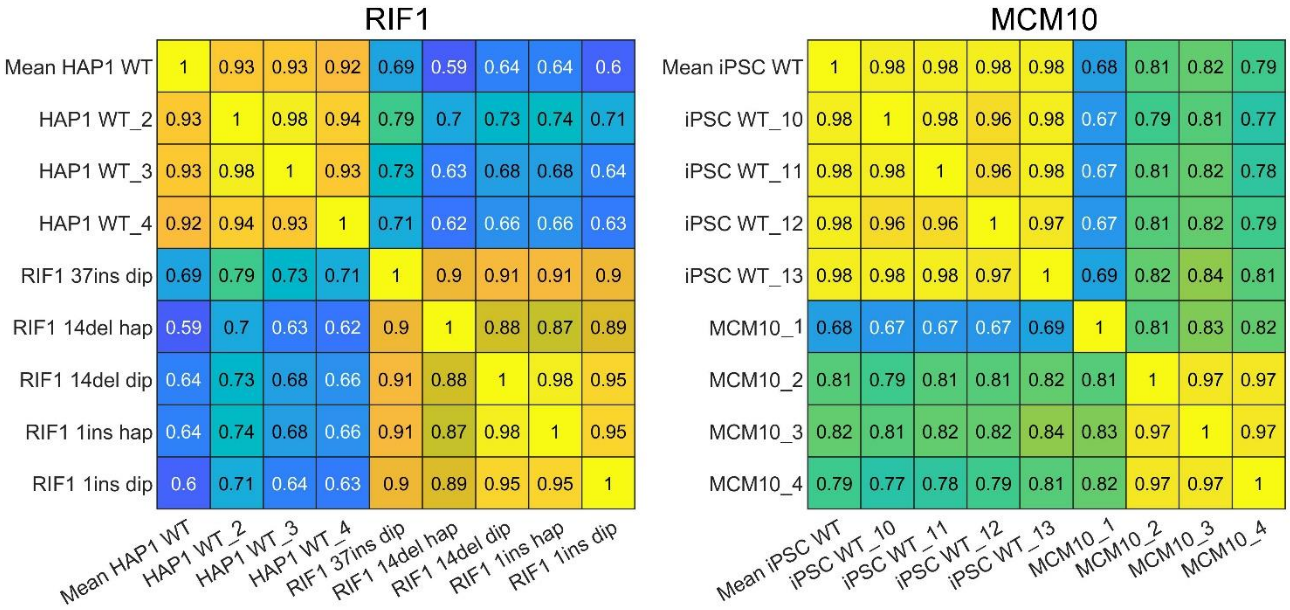


**Fig S14**. **Correlation of individual HAP1 Rif1 and iPSC MCM10 mutant samples to WT samples**.

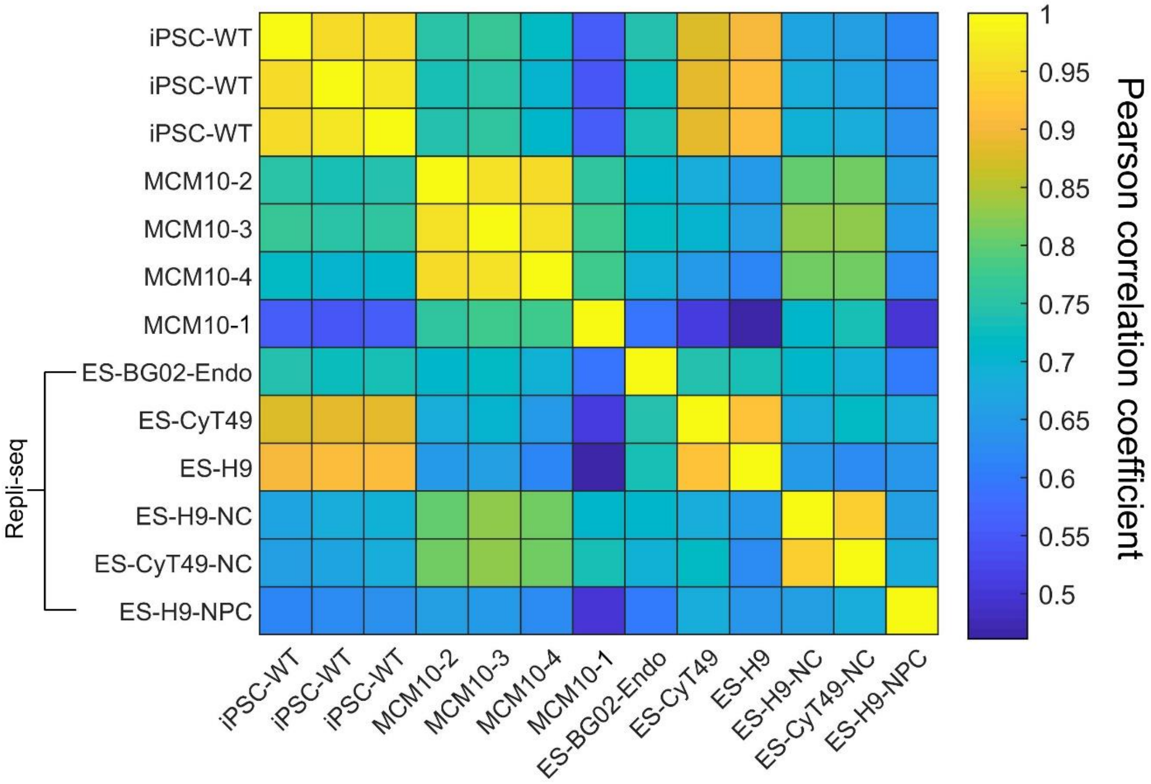


**Fig S15. MCM10 samples are not differentiated.** Whole genome correlation of replication timing profiles of select WT iPSC samples and MCM10 mutants to repli-seq profiles of ESCs and differentiated cells.

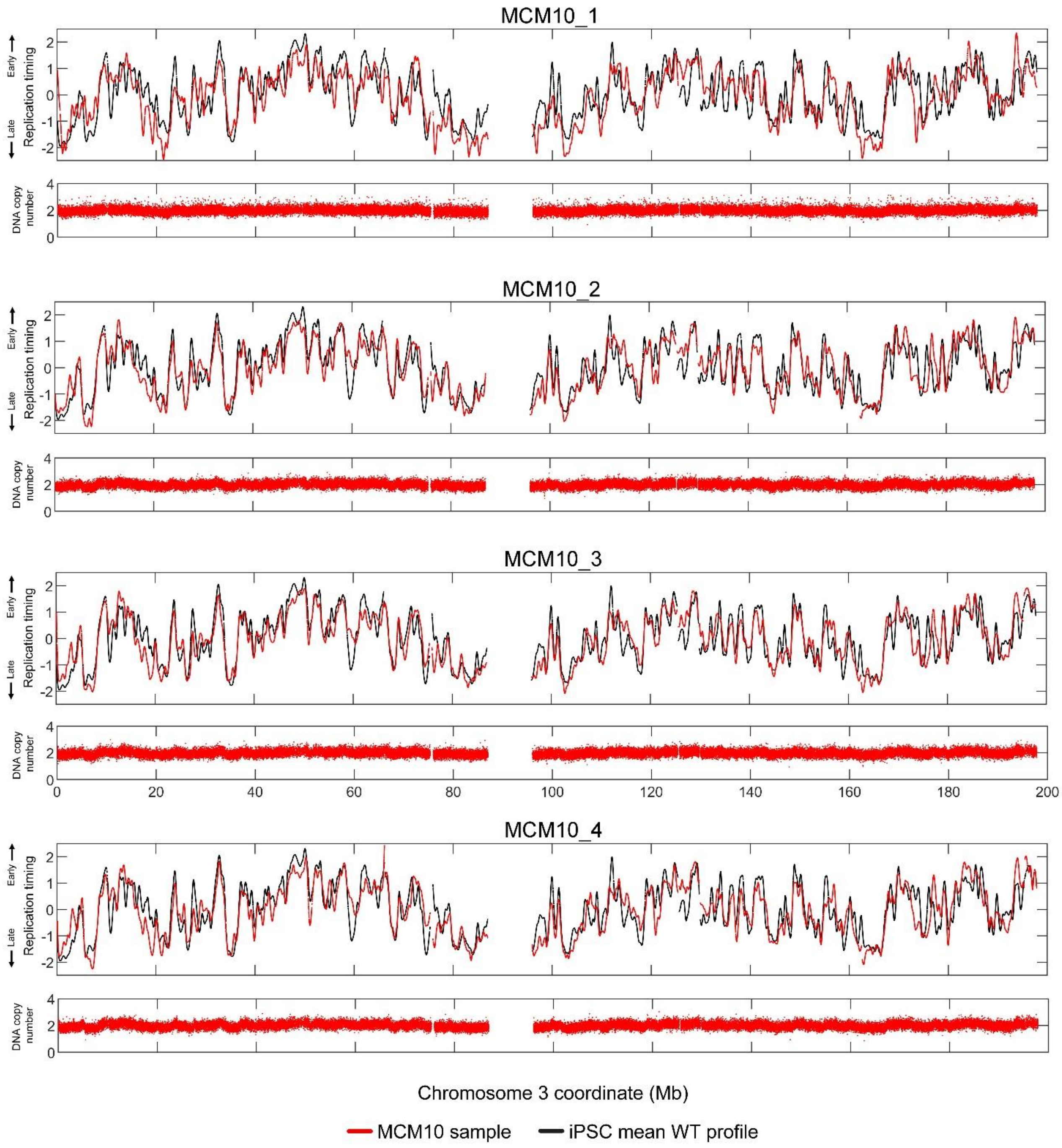


**Fig S16**. **Replication timing alterations in MCM10 are not due to copy number alterations.** The replication timing profiles (top panels, together with iPSC mean WT) and raw (pre-smoothed) copy number data (bottom panels) for each MCM10 sample. Fluctuations in DNA copy number that result from DNA replication are evident across chromosomes, however loss or gain of DNA is rare, localized and does not correspond to WT-MCM10 replication timing variation.

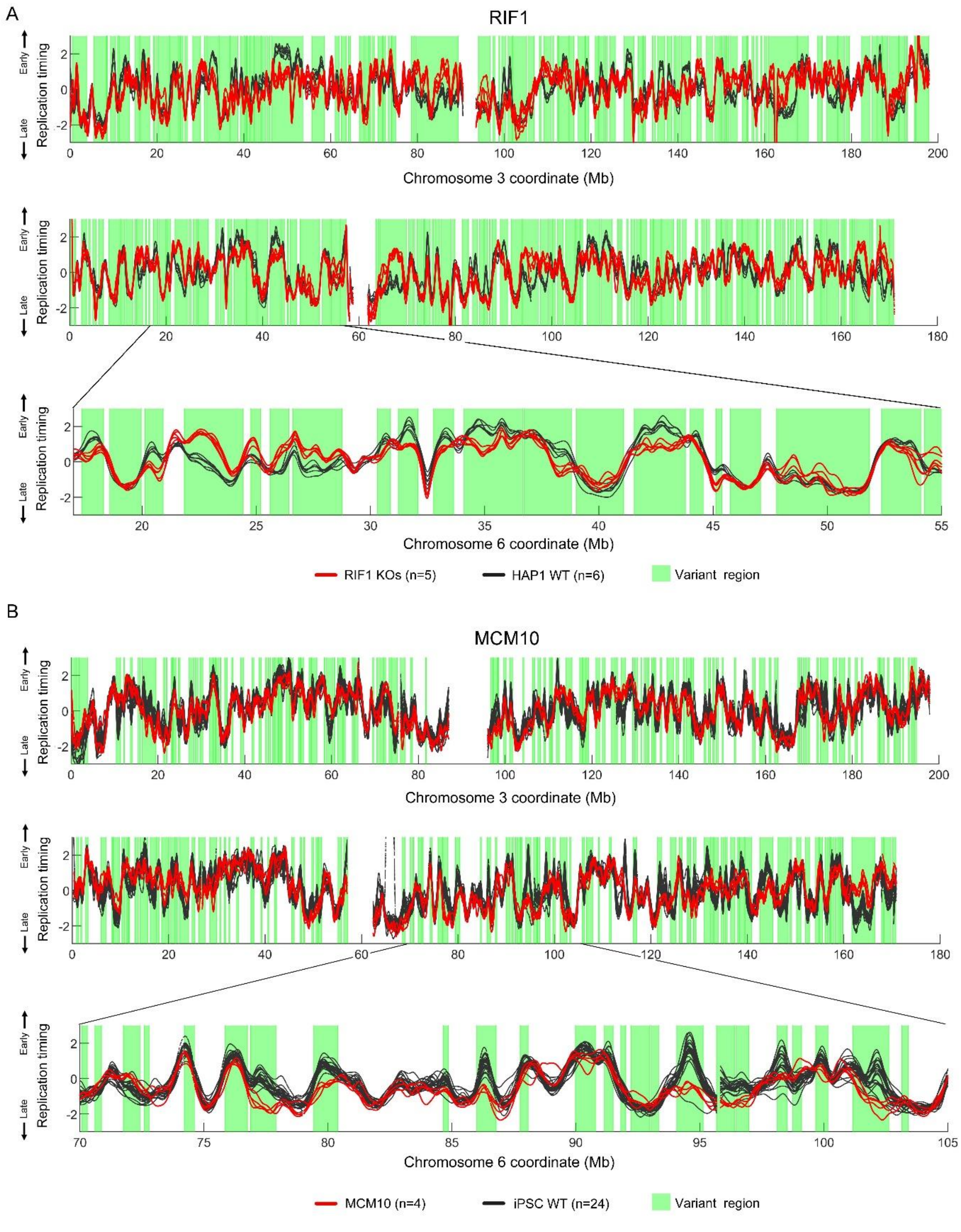


**Fig S17**. **Variation in replication timing in RIF1 and MCM10.** (A) Replication timing variant regions across RIF1 mutant samples (grouped analysis) compared to WT HAP1 samples. (B) Replication timing variant regions across MCM10 samples (grouped analysis) compared to WT iPSC samples.

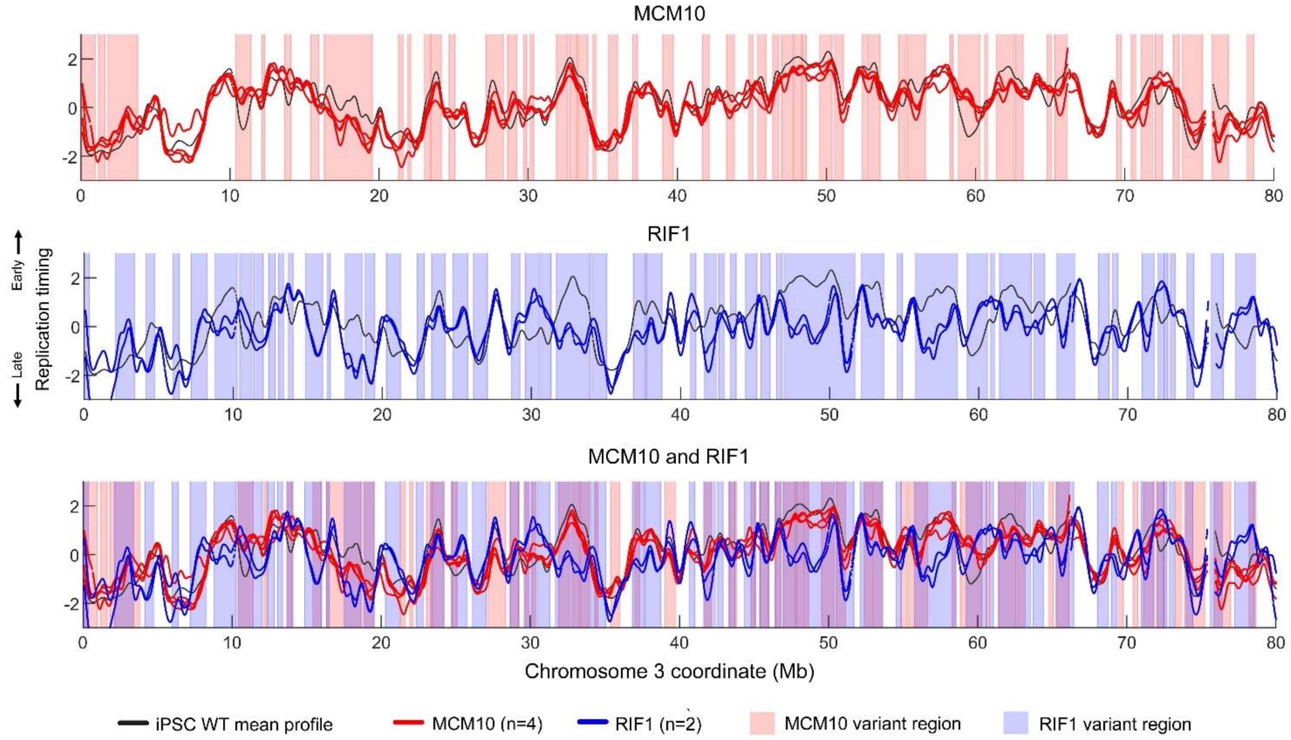


**Fig S18**. **Variation in replication timing in ESC RIF1 KO compared to MCM10.**

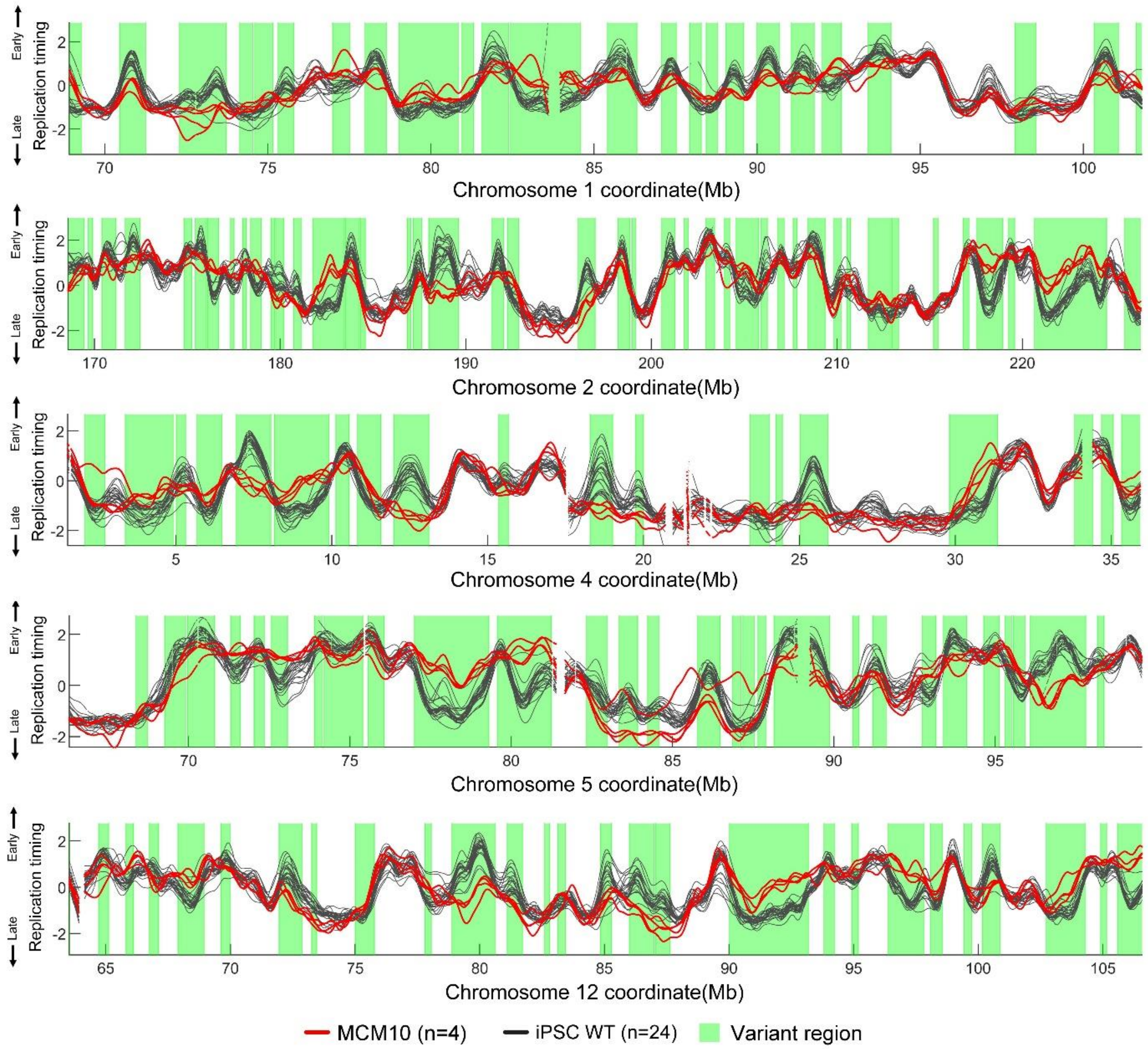


**Fig S19. Additional examples of replication timing variants in MCM10.**

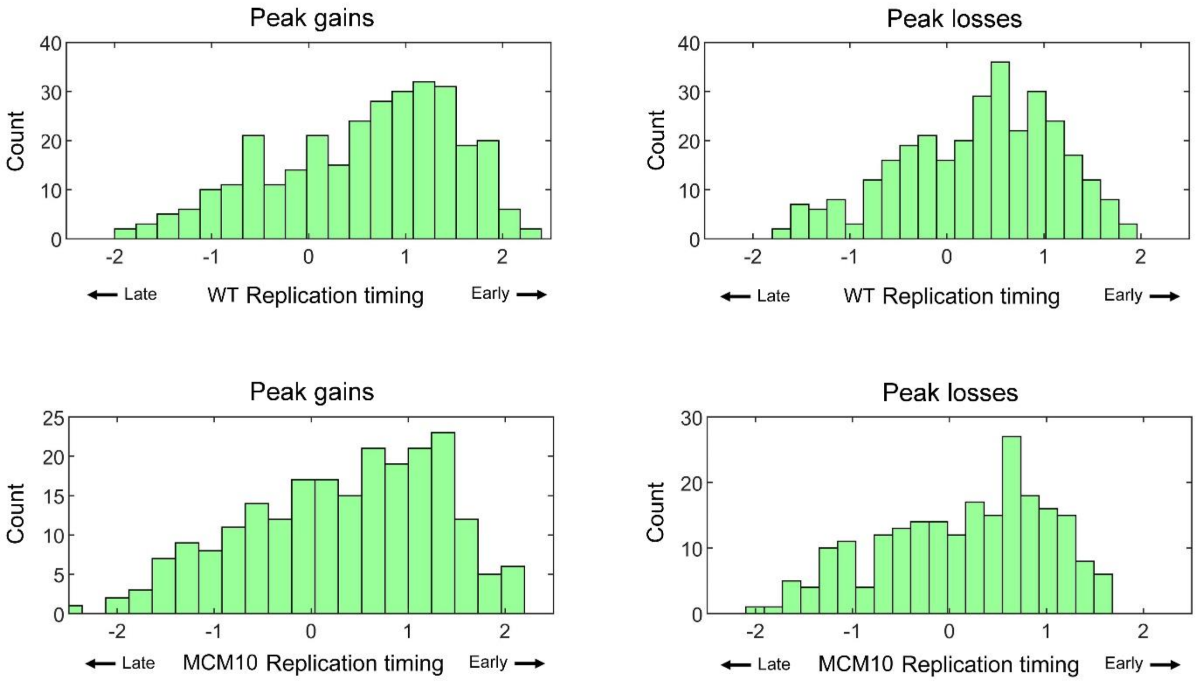


**Fig S20. The mean WT and mean MCM10 replication timing value at sites of peak gain and loss.**
